# Mono-methylated histones control PARP-1 in chromatin and transcription

**DOI:** 10.1101/2023.11.22.568326

**Authors:** Gbolahan Bamgbose, Guillaume Bordet, Niraj Lodhi, Alexei Tulin

## Abstract

PARP-1 is central to transcriptional regulation under both normal and stress conditions, with the governing mechanisms yet to be fully understood. Our biochemical and ChIP-seq-based analyses showed that PARP-1 binds specifically to active histone marks, particularly H4K20me1. We found that H4K20me1 plays a critical role in facilitating PARP-1 binding and the regulation of PARP-1-depenednt loci during both development and heat shock stress. Here we report that the sole H4K20 mono-methylase, *pr-set7*, and *parp-1 Drosophila* mutants undergo developmental arrest. RNA-seq analysis showed an absolute correlation between PR-SET7- and PARP-1-dependent loci expression, confirming co-regulation during developmental phases. PARP-1 and PR-SET7 are both essential for activating *hsp70* and other heat shock genes during heat stress, with a notable increase of H4K20me1 at their gene body. Mutating *pr-set7* disrupts monometylation of H4K20 along heat shock loci and abolish PARP-1 binding there. These data strongly suggest that H4 monometylation is a key triggering point in PARP-1 dependent processes in chromatin.

## INTRODUCTION

The chromosomes of eukaryotes are made up of chromatin, which is a complex of DNA tightly wound around nucleosomes (*1*). Nucleosomes are the basic unit of chromatin, and they are made up of about 147bp of DNA wrapped around an octamer of four core histones (H2A, H2B, H3 and H4) (*1*). Eukaryotic DNA is packed into nucleosomes to maintain higher order chromatin structure. As a consequence, DNA sequences are inaccessible to a plethora of protein effectors for biological processes such as DNA damage repair, replication and transcription (*2*). To overcome the nucleosome barrier, eukaryotes have developed several mechanisms that ensure the successful removal of nucleosomes. One of the key effectors involved in chromatin remodeling is Poly(ADP)-ribose Polymerase 1 (PARP-1), a chromatin-associated nuclear enzyme implicated in a host of biological processes including, transcriptional regulation and DNA repair (*3–5*). PARP-1 catalyzes the addition of poly(ADP)-ribose (pADPr) moieties onto acceptor proteins by utilizing donor NAD^+^ as substrate. Upon PARP-1 activation due to developmental and environmental cues or stress, PARP-1 is automodified, and it attaches negatively charged poly(ADP)-ribose polymers onto histones and other chromatin-associated proteins, thereby decondensing DNA via electro-repulsion and facilitating PARP-1 mediated biological processes (*6, 7*). Besides PARP-1, histone modifications are essential for remodeling chromatin structure. Histones are subject to post-translational modifications, including phosphorylation, acetylation, sumoylation, ubiquitylation and ADP-ribosylation on their amino-terminal tails protruding from the nucleosome core (*2, 8, 9*). Modifications of these histone tails can alter chromatin structure by modifying inter-nucleosome interactions or by serving as docking sites for effectors to facilitate diverse biological outcomes depending on the histone(s) so modified (*2, 10*).

The binding of PARP-1 to post-translationally modified histones may be necessary for PARP-1-mediated biological processes and PARP-1 targeting to chromatin. In our previous study, we showed that the phosphorylation of the histone variant H2Av directly impacts the localization, activation, and enzymatic activity of PARP-1 (*11*). Also, we previously showed that the N-terminal tails of histones H3 and H4 are needed for the enzymatic activation of PARP-1 (*12*). Taken together, there is strong evidence that histone modifications can regulate PARP-1 activity *in vivo*.

In this study, we demonstrate that PARP-1’s transcriptional regulatory role under regular and heat stress conditions may be intricately tied to its interaction with specific mono-methylated active histone marks, notably H4K20me1, H3K4me1, H3K36me1, H3K27me1 and H3K9me1 that it is inhibited by repressive H3K9me2/3 marks. Through transcriptomic analysis of *pr-set7* and *parp-1* mutant *Drosophila* third-instar larvae, we uncover a high correlation in differentially expressed genes and co-enrichment of PARP-1 and H4K20me1 at a subset of these genes, underscoring the role of PR-SET7/H4K20me1 in facilitating PARP-1-mediated transcriptional regulation. Importantly, we show that PARP-1 and PR-SET7 are essential for the activation of specific heat shock genes including *hsp70*, and the dynamic regulation of H4K20me1 during this process, further validates our hypothesis of H4K20me1 being crucial for PARP-1 binding and gene regulation under developmental and heat stress conditions.

## RESULTS

### PARP-1 binds to H4K20me1, H3K4me1, H3K36me1, H3K9me1 and H3K27me1 *in vitro* and *in vivo*

In this study, we investigated the interaction between full-length recombinant PARP-1 and various histone modifications using a histone peptide array containing modified histone peptides individually and in combination with other peptides (Fig.1A). Our previous work established that PARP-1 binds to core histones (*12*). Here, we aimed to determine the specific histone marks that modulate PARP-1’s affinity for chromatin. We found that PARP-1 binds specifically to spots containing H4K20me1, H3K4me1, H3K36me1, H3K9me1 or H3K27me1 peptides (Fig.1B, C, D and data file S1). Specificity analysis showed that PARP-1 binding was highly specific for H3K4me1, H3K36me1, H3K9me1 and H3K27me1 containing peptides with the highest specificity observed for H4K20me1 (Fig.1E and data file S1). Additionally, we explored the inhibitory effects of different histone modifications on PARP-1 binding. Our findings showed that H3K9me2/3 hindered PARP-1 binding to spots containing H3K4me1 (Fig.1F, Supplemental Fig.S1 and S2). Conversely, H3K4me1 enhanced PARP-1 binding to spots with H3K9me2/3 (Fig.1F and Supplemental Fig.S1). In parallel, our data shows that the phosphorylation of H3S10 or H3T11, two residues nearby H3K9 diminish PARP-1 binding to H3K9me1 (Supplemental Fig.S1 and S2). Similarly, the phosphorylation of H3K28 affects PARP-1 binding to H3K27me1 (Supplemental Fig.S1 and S2).

**Figure 1.**
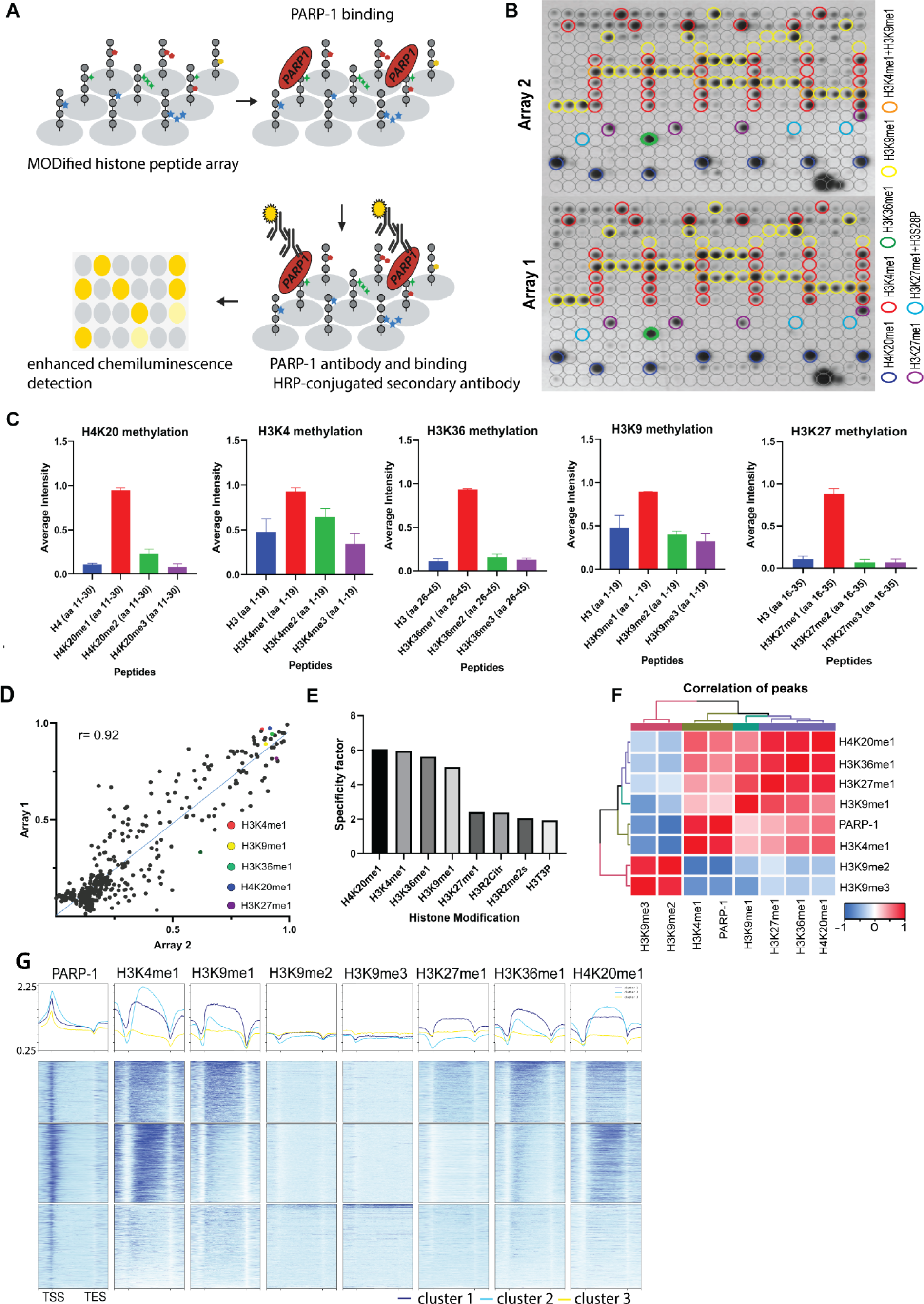
PARP-1 binds H4K20me1, H3K4me1, H3K36me1, H3K9me1, and H3K27me1 *in vitro* and *in vivo*. (**A**) Illustration of the MODified histone peptide assay (Active Motif) used to determine PARP-1 binding to histone modifications. The histone peptide array (Top, left), comprising 19mer peptides with single or up to four concurrent histone modifications, was employed to investigate PARP-1’s binding affinity for histone modifications and to assess the impact of adjacent modified peptides on PARP-1 binding. This array was first blocked, then incubated with PARP-1 protein (Top, right). Subsequently, it was stained with a PARP-1 antibody and a horseradish peroxidase (HRP)-conjugated secondary antibody (Bottom, right). Visualization of PARP-1 binding was done through enhanced chemiluminescence detection and captured on X-ray film (Bottom, left). See Methods for a full description. (**B**) Signal intensity on modified histone peptide array based on incubation with PARP-1 protein. (**C**) Average intensities of PARP-1 binding to single histone peptides. (**D-E**) Reproducibility and specificity of spot intensities from modified histone peptide array duplicates. (**D**) Scatter plot showing the correlation of the average intensities of duplicate arrays. Intensities of PARP-1 binding to all peptides and spots containing single H4K20me1, H3K4me1, H3K36me1, H3K9me1 and H3K27me1 (key) are shown. Pearson’s correlation coefficient (r) is 0.92. (**E**) Bar chart showing top 8 histone modifications with the highest specificity for PARP-1 binding. The specificity factor was calculated by dividing the average intensity of spots that contain the modified histone peptide by the average intensity of spots that do not contain the peptide. (**F**) Spearman correlation of PARP-1, H4K20me1, H3K4me1, H3K36me1, H3K9me1, H3K9me2/3 and H3K27me1 peaks in *Drosophila* third-instar larvae based on fraction of overlap. (**G**) Heatmaps showing k-means clustering-generated occupancy of PARP-1, H4K20me1, H3K4me1, H3K36me1, H3K9me1, H3K9me2/3 and H3K27me1 normalized ChIP-seq signals in third-instar larvae at *Drosophila* genes. ChIP-seq signals are sorted in descending order. The upper plots show the summary of the signals.

Next, we examined the association of PARP-1 with H4K20me1, H3K4me1, H3K36me1, H3K9me1, H3K9me2/3 and H3K27me1 in *Drosophila* third-instar larvae using ChIP-seq data. PARP-1 peaks were highly correlated with H4K20me1, H3K4me1, H3K36me1, H3K9me1 and H3K27me1 but significantly less correlated with H3K9me2/3 peaks (Fig.1G). Consistently, at gene clusters, H4K20me1 and H3K4me1 were highly enriched at gene bodies in cluster I which also had the highest PARP-1 enrichment at promoters. H3K36me1, H3K9me1 and H3K27me1 were also enriched in cluster I albeit to a lesser degree than H4K20me1 and H3K4me1 (Fig.1H). In contrast, H3K9me2/3 were more enriched at clusters II and III but depleted in cluster I (Fig.1H). Additionally, PARP-1 associates with H4K20me1 and H3K4me1 on *Drosophila* polytene chromosomes (Fig.2). Notably, H4K20me1, H3K4me1, H3K36me1, H3K9me1 and H3K27me1 are associated with active genes while H3K9me2/3 is associated with low-expression or silent genes (*13*). Thus, our data shows that PARP-1 predominantly binds to active mono-methylated histone marks, specifically H4K20me1, H3K4me1, H3K36me1, H3K9me1 and H3K27me1 while also suggesting that the binding of PARP-1 may be hindered by repressive histone marks such as H3K9me2/3.

**Figure 2.**
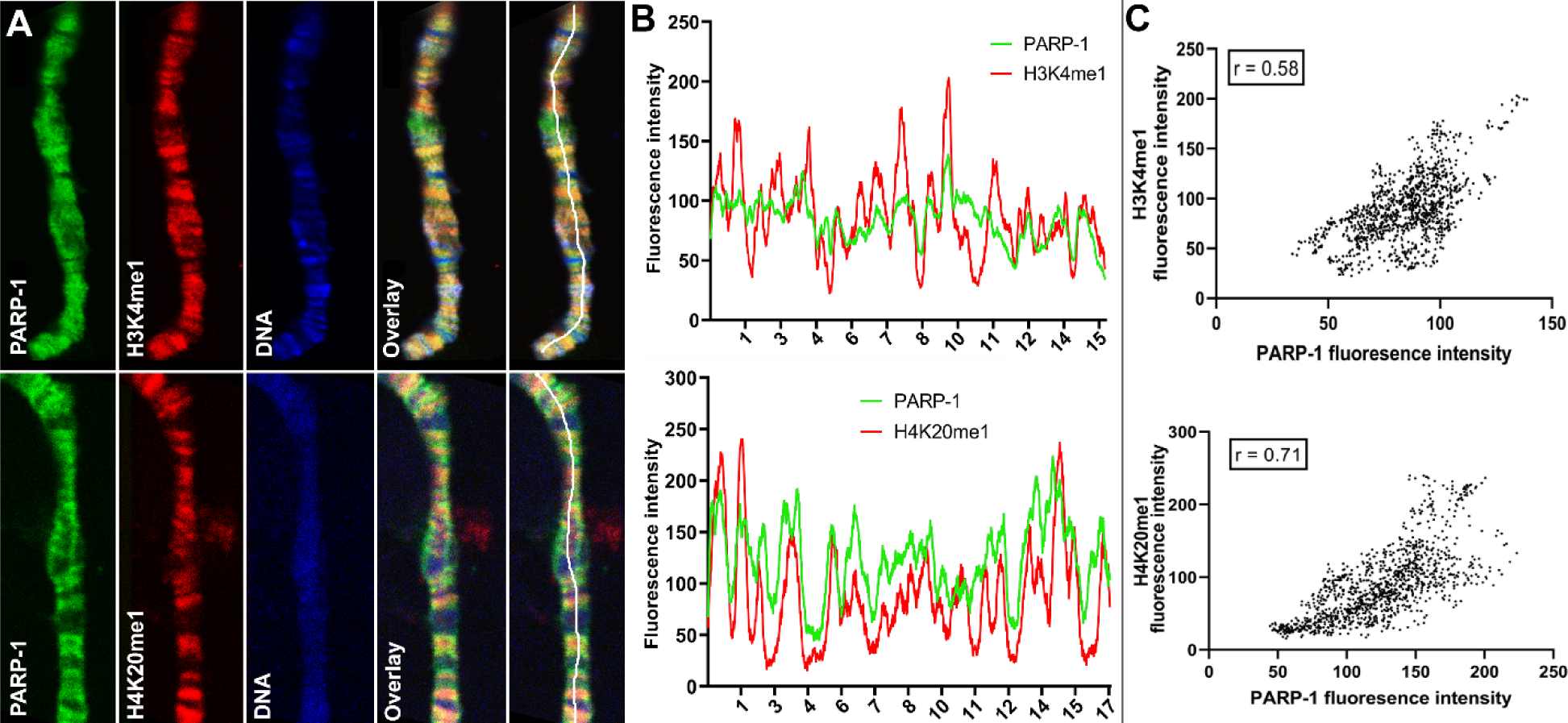
PARP-1 colocalizes with H3K4me1 and H4K20me1 histone marks in polytene chromosomes. (**A**) Immunofluorescent staining of *Drosophila* salivary gland chromosomes showing colocalization of PARP-1 with H3K4me1 and H4K20me1. White lines indicate areas of the colocalization-quantification shown in (**B**). (**B-C**) Quantification of fluorescence intensity of PARP-1, H3K4me1 and H4K20me1 at *Drosophila* polytene chromosomes in panel A. (**B**) Images show distribution of PARP-1 fluorescence intensity with H3K4me1 (top) and H4K20me1 (bottom) fluorescence intensity. (**C**) Images represent a scatterplot showing PARP-1 fluorescence intensity with H3K4me1 (top) and H4K20me1 (bottom) fluorescence intensity. Pearson correlation coefficients (r) are respectively 0.58 (top) and 0.71 (bottom).

### PARP-1 and PR-SET7 co-regulate developmental gene expression programs

Next, we examined the relationship between H4K20me1 and PARP-1 during transcriptional regulation. Given that flies that lack PR-SET7, the sole methylase of H4K20, and PARP-1 undergo developmental arrest during the larval-to-pupal transition (*14, 15*), we hypothesized that PR-SET7/H4K20me1 and PARP-1 may regulate similar gene expression programs during development. To validate our hypothesis, we initially confirmed that the *pr-set7^20^*mutant not only eliminated PR-SET7 RNA and protein but also abrogated H4K20me1 modification (Supplemental Fig.S3). Interestingly, in the absence of PARP-1, neither PR-SET7 RNA nor protein levels were affected (Supplemental Fig.S4-5), indicating that PARP-1 is not directly implicated in the regulation of *pr-set7*. This finding contrasts with recent evidence demonstrating PARP1-induced degradation of PR-SET7/SET8 in human cells (*16*). Subsequently, we integrated PARP-1 and H4K20me1 ChIP-seq data in third-instar larvae and RNA-seq of *pr-set7^20^* and *parp-1^C03256^* mutant third-instar larvae. Our data showed that the differentially expressed genes (DEGs) in *pr-set7^20^* and *parp-1^C03256^*transcriptome were highly correlated (Pearson, r = 0.79) (Fig.3A and B). However, despite the global reduction of H4K20me1, and PARP-1 RNA in *pr-set7^20^*and *parp-1^C03256^* mutants, respectively (*14, 15*), there was a limited alteration in gene expression at genes where PARP-1 was co-localized with H4K20me1 (Fig.3C and D). Differentially expressed genes in both *pr-set7^20^* and *parp-1^C03256^*mutants and co-enriched with PARP-1 and H4K20me1 were mainly upregulated (n = 101, 72%) (Fig.3E). Intriguingly, under wild-type conditions, these genes displayed expression levels approximately 40% higher than the average and demonstrated increased of RNA-Polymerase II occupancy both at their promoter regions and gene bodies compared to other genes (Supplemental Fig.S6), indicating their high activity in wild type context. Interestingly, among the genes co-enriched with PARP-1 and H4K20me1, those only differentially expressed in *parp-1^C03256^* mutants were mostly upregulated (n = 94, 85%), while those only differentially expressed in *pr-set7^20^* mutants were mainly downregulated (n = 191, 67%), indicating that PARP-1 and H4K20me1 also have distinct functions in gene regulation at co-enriched genes (Fig.3E). Gene ontology analysis of differentially expressed genes bound by PARP-1 and H4K20me1 showed that metabolic genes were upregulated while genes involved in neuron development and morphogenesis were downregulated in both *pr-set7^20^*and *parp-1^C03256^* mutants (Fig.3F). Finally, to extend the generalizability of our observations beyond *Drosophila*, we compared the distribution of PARP1 and H4K20me1 in Human K562 cells. Strikingly, we observed a correlation in their distribution, suggesting that the interplay between PARP-1 and H4K20me1 is not limited to fruit flies (Supplemental Fig.S7). Overall, our results indicate H4K20me1 may be required for PARP-1 binding to preferentially repress metabolic genes and activate genes involved in neuron development at co-enriched genes.

**Figure 3.**
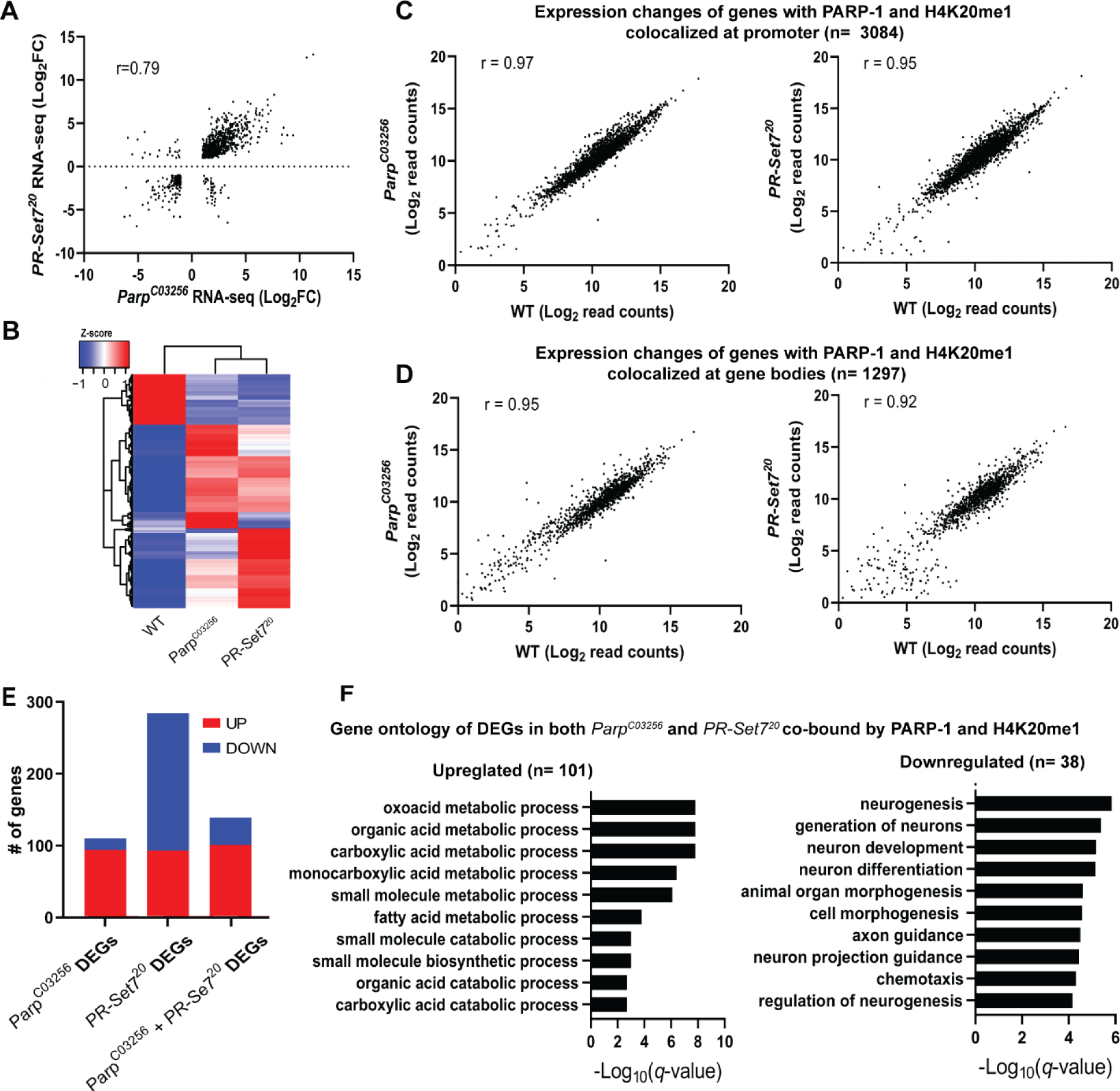
PARP-1 and H4K20me1 are required for the repression of metabolic genes and activation of developmental genes at co-enriched genes. (**A**) Scatterplot plot showing correlation of differentially expressed genes (DEGs) in *parp-1^C03256^*and *pr-set7^20^* (Pearson’s r = 0.79). (**B**) Heatmap showing the normalized read counts of DEGs in both *parp-1^C03256^* and *pr-set7^20^*. Normalized read counts are shown as row z-scores. (**C-D**) Dot plots showing transcriptional changes of genes co-enriched with PARP-1 and H4K20me1 in *parp-1^C03256^* and *pr-set7^20^*compared to WT at promoters (**C**) and gene bodies (**D**). (**E**) Summary of DEGs in *parp-1^C03256^* and *pr-set7^20^* and both mutants that were co-enriched with PARP-1 and H4K20me1. (**F**) Gene ontology of upregulated (left) and downregulated (right) DEGs in both *parp-1^C03256^* and *pr-set7^20^* mutants that were co-enriched with PARP-1 and H4K20me1.

### PARP-1 and PR-SET7/H4K20me1 are required for the optimal expression of heat shock genes during heat stress

Next, we examined PARP-1 and H4K20me1 controlled gene expression programs during dynamic transcriptional changes. During heat shock, PARP-1 spreading from the promoter to the gene bodies of *hsp70* facilitates nucleosomal loss which leads to transcriptional activation (*17, 18*). Since H4K20me1 is mostly enriched in gene bodies (Fig.1H), we hypothesized that H4K20me1 may be required for PARP-1 spread at *hsp70* during heat shock. First, we examined the expression of heat shock genes before and after 30 minutes heat shock in WT, *parp-1^C03256^ and pr-set7^20^*third-instar larvae. The expression of heat shock genes (*hsp22, hsp23, hsp68, hsp83*) including *hsp70* were significantly reduced in *parp-1^C03256^ and pr-set7^20^* animals after heat shock (Fig.4A). ChIP-seq results revealed that PARP-1 highly occupied the promoters of *hsp23*, *hsp68*, *hsp70*, and *hsp83*, as well as in the gene body of *hsp22*, prior to heat shock (Fig. 4B and C). Analysis of H4K20me1 enrichment showed low levels across the gene bodies of *hsp22, hsp68*, and *hsp70* (group A) but high levels at gene bodies of *hsp23* and *hsp83* (group B) before heat shock (Fig.4B and C). Based on these findings, we hypothesized that an increase in H4K20me1 enrichment in the gene bodies of group A genes would facilitate PARP-1 binding after heat shock, while H4K20me1 enrichment remains unchanged in group B genes thereby facilitating PARP-1 binding and spreading. To our surprise, H4K20me1 levels increased at the gene bodies of group A genes (low H4K20me1 before heat shock) (Fig.4D) but decreased significantly at group B genes (high H4K20me1 before heat shock) after heat shock (Fig.4E). These findings suggest that dynamic changes in H4K20me1 enrichment may regulate the expression of heat shock genes during heat shock and indicate that H4K20me1 may not be necessary for PARP-1 binding and PARP-1-mediated activation of group B genes during heat shock.

**Figure 4.**
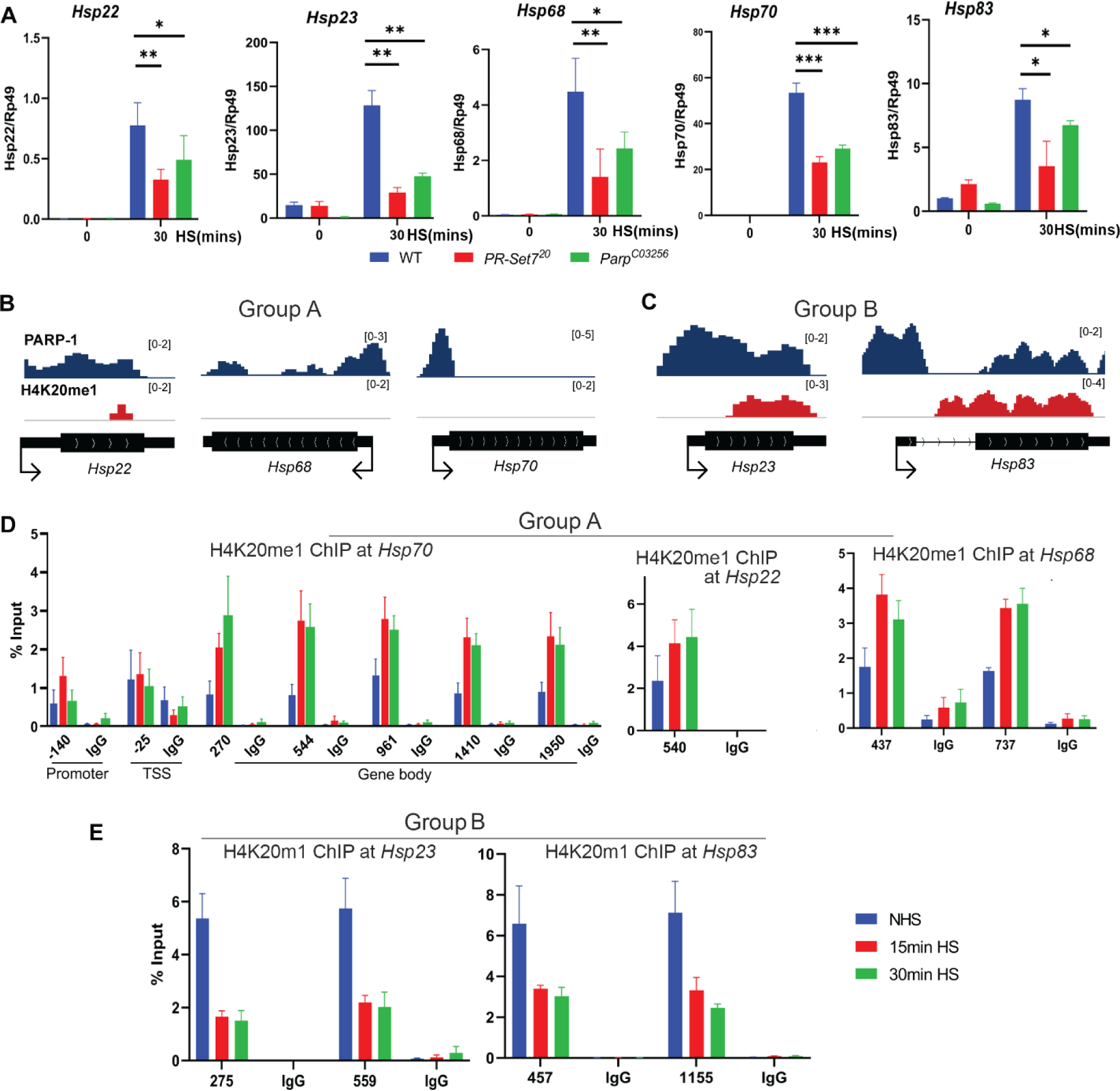
Dynamic H4K20me1 enrichment regulates the expression of heat-shock genes during heat shock. (**A**) Expression of *hsp22, hsp23, hsp68, hsp70* and *hsp83* in WT, *parp-1^C03256^* and *pr-set7^20^*third-instar larvae before and after 30 mins heat shock. Data shown are from 3-5 biological replicates. \*\*\**P* < 0.001, \*\**P* < 0.01, \**P* < 0.05 (Unpaired t-test; Two-tailed). Integrative Genome Viewer (IGV) tracks showing normalized ChIP-seq tracks of PARP-1 and H4K20me1 in third-instar larvae at (**B**) *hsp22, hsp68, hsp68, hsp70* and (**C**) *hsp22, hsp83* before heat shock. Mononucleosome ChIP-qPCR in WT showing enrichment of H4K20me1 at (**D**) *hsp70, hsp22, hsp68* and (**E**) *hsp23* and *hsp83* before heat shock (NHS), after 15 mins heat shock and 30 mins heat shock. Primers used spanned the *hsp70* locus and the gene bodies of *hsp22, hsp23, hsp68, hsp70* and *hsp83*. Data shown are from 3 biological replicates.

### Mutating PR-SET7 methyl transferase disrupts PARP-1 binding to chromatin and diminishes poly(ADP)-ribosylation levels during heat shock stress

Our results showed that prior to heat shock, there was no effect on PARP-1 binding in both WT and *pr-set7^20^* background animals to chromatin (Fig.5A). However, after heat shock, we observed that PARP-1 binding was concentrated at specific loci in WT animals, while it was diminished from chromatin in *pr-set7^20^*animals (Fig.5A). These findings suggest that PR-SET7/H4K20me1 plays a crucial role in mediating PARP1 binding to chromatin during heat stress, thus, controls PARP1-dependent processes in chromatin. Previous studies showed that heat shock results in pADPr accumulation at the heat shock-induced puffs during PARP-1-dependent transcriptional activation (*5, 7, 11, 17–19*). Therefore, we tested if heat-shock treatment alters pADPr accumulation in *pr-set7^20^*animals. As expected, the pADPr level increased by 26.1 times after heat-shock treatment in wild type animals, while in *pr-set7^20^* mutants the level of pADPr increased only 2.3 folds (Fig.5B). This observation implies that PR-SET7/H4K20me1 plays a key role in regulating PARP-1 upon environmental stresses such as heat shock.

**Figure 5.**
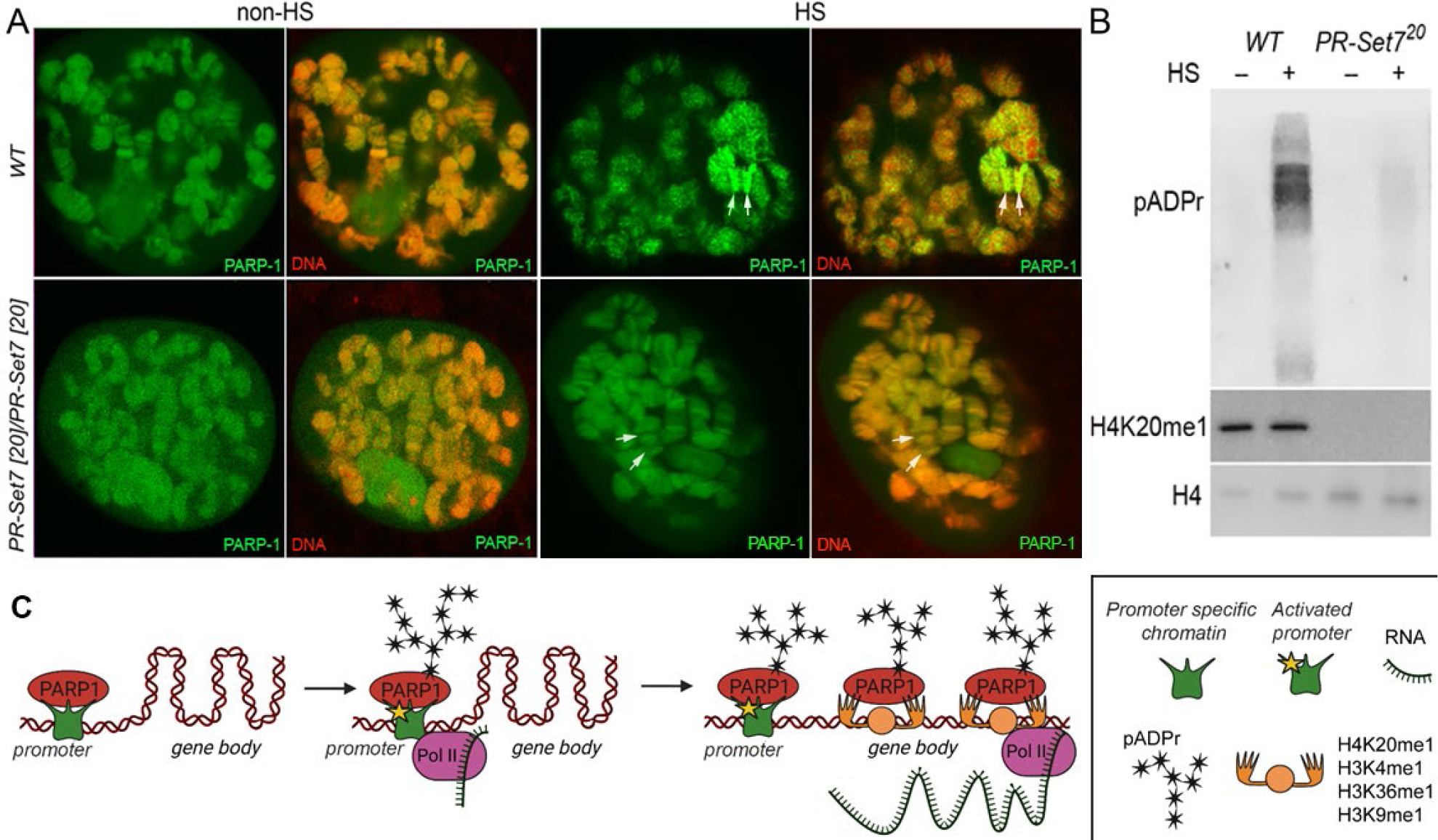
Mono-methylated histones controls PARP-1 binding along gene body to regulate transcription. (**A**) PR-SET7/H4K20me1 is required for holding PARP-1 in chromatin during heat shock (HS). PARP-1 (green) protein recruitment to *hsp70* locus (Arrows) in salivary gland polytene nuclei in wild type (WT) and *pr-set7^20^* mutant third instar larvae. A single salivary gland polytenized nucleus of wandering third instar larvae is shown for each genotype. Wild type genotype - UASt::PARP-1-EYFP, GAL4[Mz1087.hx]; *pr-set7* mutant genotype – UASt::PARP-1-EYFP, GAL4^Mz1087.hx^; *pr-set7^20^*. (**B**) Equal amounts of lysates from the wild-type expressing PARP-1-EYFP (WT) and *pr-set7^20^*mutants expressing PARP-1-EYFP (*pr-set7^20^*) grown at 22°C or heat shocked at 37°C for one hour at the third-instar larvae stage were subjected to immunoblot analysis using mouse anti-pADPr (10H), anti-H4K20me1 and anti-histone H4 (loading control) antibodies. (**C**) **Model of PARP-1 regulation by histone modifications.** PARP-1 binds to a nucleosome that carries the H2A variant (H2Av) at the promoter region. Upon developmental triggers or heat shock-induced phosphorylation of H2Av, PARP-1 is activated (*11*). Activated PARP-1 fosters a transcription start site (TSS) that is more accessible, thereby enabling the binding of RNA-Polymerase II (Pol II) and initiation of transcription (*26*). Following this, the distribution of PARP-1 is further enhanced by active mono-methylated forms of H4K20, H3K4, H3K36, or H3K9. Each of these histone modifications can interchangeably facilitate this function. The spreading of PARP-1 to the gene body contributes to the loosening of chromatin in this region (*7*, *16*, *17*, *49*). Consequently, this facilitates the transition of Pol II into a productive elongation phase, leading to the generation of a mature transcript.

## DISCUSSION

This study illuminates the intricate interactions between PARP-1 and diverse histone modifications, with a particular emphasis on the influence of histone marks on the chromatin binding propensity of PARP-1. We discovered an elevated binding affinity of PARP-1 towards specific mono-methylated active histone modifications such as H4K20me1, H3K4me1, H3K36me1, H3K9me1 and H3K27me1 *in vitro* (Fig.1B and C). Intriguingly, our results also suggest that the repressive histone modifications, H3K9me2/3 may potentially hinder PARP-1 binding (Fig.1F). The association between PARP-1 peaks and distinct histone marks in *Drosophila* third-instar larvae, as revealed through ChIP-seq data, corroborates the significance of these histone marks in PARP-1 binding. This is predominantly apparent for H4K20me1, H3K4me1, H3K36me1, H3K9me1 and H3K27me1, while H3K9me2/3 demonstrated a lesser correlation (Fig.1G), further validating the repressive role of these marks.

We additionally examined the potential functional consequences of PARP-1’s interaction with these histone marks, particularly H4K20me1. Mono-methylation of histone H4 on lysine 20 (H4K20me1), catalyzed by PR-SET7, is evolutionarily conserved from yeast to humans (*20, 21*). H4K20me1, similar to PARP-1, is enriched at highly active genes (*22–28*). Also, H4K20me1 is critical for transcriptional regulation, maintenance of genome integrity, cell cycle regulation and double-strand break (DSB) DNA damage repair, all biological processes in which PARP-1 plays a role (*3, 20, 29–33*). Hence, histone modifications such as H4K20me1 may act to regulate the activity of key chromatin remodelers such as PARP-1.

A recent study demonstrated that in human cells overexpressing PARP-1, PR-SET7/SET8 is degraded, whereas depletion of PARP-1 leads to an increase in PR-SET7/SET8 levels (*16*). However, in our study involving *parp-1* mutant in *Drosophila* third-instar larvae revealed a nuanced scenario: we detected a minor but not significant reduction in both PR-SET7 RNA and protein levels (Supplemental Fig.S4 and S5). This outcome stands in stark contrast to the previous study’s findings. The discrepancy could be due to the distinct experimental approaches used: the previous research focused on mammalian cells and *in vitro* experiments, whereas our study examined the functions of PARP-1 in whole *Drosophila* third-instar larvae during development. Consequently, while PARP-1 may cooperate with PR-SET7 in the context of *Drosophila* development, it could exhibit antagonistic roles against PR-SET7 in specific cell lines and under certain biological or developmental conditions.

Our transcriptomic analysis from *pr-set7^20^* and *parp-1^C03256^* mutant third-instar larvae implies that PR-SET7/H4K20me1 might be crucial for PARP-1’s role in transcriptional regulation. Remarkably, differentially expressed genes (DEGs) in *pr-set7^20^*and *parp-1^C03256^* mutants exhibited a high degree of correlation (Fig.3A), although we noticed minimal alteration in gene expression where PARP-1 was co-localized with H4K20me1 across the genome (Fig.3C and D). This suggests that H4K20me1 might only be essential for PARP-1-mediated binding and gene regulation at a select group of genes, specifically during metabolic gene repression and developmental gene activation. Our results also propose that PARP-1 and PR-SET7/H4K20me1 might have separate roles in gene regulation at co-enriched genes.

Notably, genes co-enriched with PARP-1 and H4K20me1, and are upregulated in both *parp-1^C03256^* and *pr-set7^20^*mutants, are predominantly metabolic genes exhibiting high expression levels under wild-type conditions and a high occupancy of RNA-polymerase II both at their promoter region and gene body (Supplemental Fig.S6). In our previous studies, we discovered that PARP-1 plays a crucial role in repressing highly active metabolic genes during the development of *Drosophila* by binding directly to their loci (*34, 35*). Also, PARP-1 is required for maintaining optimum glucose and ATP levels at the third-instar larval stage (*34*). During *Drosophila* development, repression of metabolic genes is crucial for larval to pupal transition (*36, 37*). This repression is linked to the reduced energy requirements as the organism prepares for its sedentary pupal stage (*36, 38*). Remarkably, our observations indicate a notable affinity of PARP-1 for binding to the gene bodies of these metabolic genes (*34*), suggesting a direct involvement of PARP1 in their regulation. Nonetheless, it remains plausible that certain genes may be indirectly regulated by PARP1 through intermediary transcription factors.

Our data indicates that in both *parp-1* and *pr-set7* mutant animals, there was a preferential repression of metabolic genes at sites where PARP-1 and H4K20me1 are co-bound (Fig.3E), while these metabolic genes are highly active during third-instar larval stage (Supplemental Fig.S6). Thus, we propose that the presence of H4K20me1 may be essential for the binding of PARP-1 at these gene bodies, contributing to their repression. Importantly, this mechanism of gene repression has broader developmental implications. As earlier stated, mutant animals lacking functional PARP-1 and PR-SET7 undergo developmental arrest during larval to pupal transition. This arrest could be directly linked to the disruption of the normal metabolic gene repression during development. Without the repressive action of PARP-1 and PR-SET7, key metabolic processes might remain unchecked, leading to metabolic imbalances that are incompatible with the normal progression to the pupal stage.

In this study, we have demonstrated that PARP-1 is essential for the activation of *hsp22, hsp23,* and *hsp83* during heat shock (Fig.4A). Importantly, the expression of these heat shock genes significantly decreased in *pr-set7^20^* mutants following heat shock (Fig.4A), which shows a role for PR-SET7/H4K20me1 in the heat shock response. We have also observed that H4K20me1 regulation is dynamic in the gene bodies of PARP-1- and PR-SET7-dependent heat shock genes. Specifically, at genes that show low or no H4K20me1 enrichment prior to heat shock, H4K20me1 levels increase post-heat shock (Fig.4B and D). Conversely, at genes with high levels of H4K20me1, the level of H4K20me1 decreases following heat shock (Fig.4C and D). Previous studies have shown that H4K20me1 occupancy can either inhibit or facilitate transcriptional elongation, depending on the biological context (*26, 28, 39–41*). As such, we propose that for group A genes, which show a significant increase in H4K20me1 enrichment after heat shock, H4K20me1 may facilitate the spread of PARP-1 in their gene bodies, thus enhancing transcriptional activation. In contrast, the removal of H4K20me1 at group B genes, which exhibit a significant reduction of H4K20me1 at their gene bodies, may enable transcriptional elongation. Thus, an unidentified H4K20me1 demethylase and PR-SET7’s non-catalytic functions may be required to facilitate transcriptional activation of group B heat shock genes during heat stress.

Another plausible explanation could be that the recruitment of PARP-1 to group B genes loci promotes H4 dissociation and then leads to a reduction of H4K20me1. However, our findings suggest an alternative interpretation: the decrease in H4K20me1 at group B genes during heat shock does not seem to impede PARP-1’s role in transcriptional activation, (Fig.4A, C and E). Rather than disrupting PARP-1 function, we propose that this reduction in H4K20me1 may signify a regulatory shift in chromatin structure, priming these genes for transcriptional activation during heat shock, with PARP-1 playing an independent facilitating role. Moreover, existing studies have highlighted the dual role of H4K20me1, acting as a promoter of transcription elongation in certain contexts and as a repressor in others (*13, 26, 39, 40, 42–46*). The elevated enrichment of H4K20me1 in group B genes under normal conditions may indicate a repressive state that requires alleviation for transcriptional activation. Additionally, we cannot discount the possibility of unique regulatory functions associated with PR-SET7, extending beyond its recognized role as a histone methylase. Non-catalytic activities and potential interactions with non-histone substrates might contribute to the nuanced control exerted by PR-SET7 on group B genes during heat stress (*47, 48*). Furthermore, our exploration of *pr-set7^20^*and *parp-1^C03256^* mutants reveals distinct roles for PARP-1 and H4K20me1 in modulating gene expression (Fig 3E). This reinforces the notion that the interplay between PR-SET7 and PARP-1 involves a multifaceted regulatory mechanism. Understanding the intricate relationship between these molecular players is crucial for elucidating the complexities of gene expression modulation under heat stress conditions.

Notably, *Drosophila* embryo expressing an unmodifiable H4K20 (H4K20A) and a catalytically-inactive *trr* mutant (monomethylase of H3K4) were still able to mature into adult flies (*14, 49*), demonstrating that H3K4me1 and H4K20me1 are dispensable for transcription during development. This finding contrasts our hypothesis that H4K20me1 or H3K4me1 might modulate PARP-1 binding at PARP-1-dependent genes during development. Importantly, our data also revealed that PARP-1 binds to H3K36me1 and H3K9me1. Consequently, we suggest that H3K36me1 and H3K9me1 could potentially substitute the role of H3K4me1 and H4K20me1 in controlling PARP-1 binding at PARP-1 dependent genes (Fig.5C).

Finaly, highly transcribed genes have been reported to present a high turnover of mono-methylated modifications, maintaining a state of low methylation (*50*). Moreover, our previous study revealed that PARP1 preferentially binds to highly active genes (*34*). Consequently, our findings suggest an active involvement of PARP-1 in the turnover process to maintain an active chromatin environment. This proposed mechanism unfolds in the following steps: 1) PARP-1 selectively binds to mono-methylated active histone marks associated with highly transcribed genes. 2) Upon activation, PARP-1 undergoes automodification and subsequently disengages from chromatin, facilitating the reassembly of nucleosomes carrying the mono-methylated marks. 3) The enzymatic action of Poly(ADP)-ribose glycohydrolase (PARG) cleaves pADPr, restoring PARP-1’s binding affinity to mono-methylated active histone marks. This proposed hypothesis is consistent with existing research conducted across various model organisms, including mice, Drosophila, and Humans (*7, 24, 30, 51–53*). Notably, previous studies have consistently demonstrated that PARP-1 predominantly associates with highly expressed genes and plays a crucial role in mediating nucleosome dynamics and assembly. Thus, our proposed model provides a molecular framework that may contribute to understanding the relationship between PARP-1 and the epigenetic regulation of gene expression.

## MATERIALS AND METHODS

### *Drosophila* strains and genetics

Flies were cultured on standard cornmeal-molasses-agar media at 25°C. The transgenic stock pP{w^1^, UAST::PARP1-EYFP} has been previously described (*52*). The *parp-1^C03256^*hypomorph mutant was generated in a single pBac-element mutagenesis screen (*54*). *Parp-1^C03256^* significantly lowers the level of PARP-1 RNA and protein level (*15*) but also significantly diminishes the level of pADPr (*11*). The *pr-set7^20^* null mutant was generously provided by Dr. Ruth Steward (*14*). The *pr-set7^20^* null mutant was validated in (*14*) and we confirmed that this mutant abolishes PR-SET7 RNA and protein level but also leads to the absence of H4K20me1 (Supplemental Fig.S3). Wandering third-instar larvae was used for all experiments. *w^1118^* strains were used as controls for ChIP-seq and RNA-seq and termed WT as appropriate. We used TM6B and TM3-GFP to isolate *parp-1^C03256^* and *pr-set7^20^* homozygous mutants respectively.

### Histone peptide array

The Modified Histone peptide array (Active Motif) was blocked by overnight incubation in the tween 20 containing tris buffered saline (TTBS) buffer (10 mm Tris/HCl pH 8.3, 0.05% Tween-20 and 150 mm NaCl) containing 5% non-fat dried milk at 4°C. The membrane was then washed once with the TTBS buffer and incubated with 4.0μgm PARP-1 (Trevigen) in PARP binding buffer (10 mm Tris-HCl, pH 8, 140 mm NaCl, 3 mm DTT, and 0.1% Triton X-100) at room temperature for 1 h. The membrane was washed in the TTBS buffer and incubated with anti-PARP antibody (SERUTEC at 1:500 dilution) for 1 h at room temperature in blocking buffer (5% milk in TTBS). The unbound antibody was washed three times with TTBS, and the membrane was incubated with horseradish peroxidase-conjugated anti-mouse antibody (Sigma, 1:2500) in TTBS for 1 h at room temperature. Finally, the membrane was submerged in ECL developing solution (GE Healthcare) and the image was captured on X-ray film. Typical exposure times were 0.5–2 min. The images were analyzed using an in-house program (Array Analyze, available at https://www.activemotif.com/catalog/668/modified-histone-peptide-array).

### *Drosophila* salivary gland polytene chromosome immunostaining

Preparation and immunostaining of polytene chromosome squashes were performed exactly as described (*55*). The primary antibody used was anti-GFP (Living Colors, #JL-8, 1:100), H3K4me1 (Abcam, 1:50) and H4K20me1 (Abcam, 1:100) and the secondary antibody used was goat anti-mouse Alexa-488 (Molecular Probes, 1:1500) and goat-anti-rabbit Alexa-568 (Molecular Probes, 1:1500). DNA was stained with TOTO3 (1:3000). Slides were mounted in Vectashield (Vector Laboratories, Burlingame, CA).

### Chromatin immunoprecipitation and sequencing

PARP-1-YFP ChIP-seq has been previously described (*35*). Briefly, 75 third-instar larvae were collected, washed with 1ml 1X PBS, and homogenized in lysis buffer (200ul 1X protease inhibitor cocktail, 250ul PMSF, 800ul 1X PBS, 1ul Tween 20). Crosslinking was achieved by adding 244.5ul of 11% formaldehyde for a final concentration of 1.8% and quenched with 500mM glycine. The pellet was resuspended in sonication buffer (0.5% SDS, 20mM Tris pH 8.0, 2mM EDTA, 0.5mM EGTA, 0.5mM PMSF, 1X protease inhibitor cocktail) and sonicated to fragment chromatin. The supernatant was then collected after pelleting. This was pre-cleared, incubated with anti-GFP antibody (TP-401, Torrey Pines Biolabs), and immunoprecipitated chromatin was collected using Protein A agarose beads. Sequential washing was performed using low salt buffer, high salt buffer, LiCl wash, and TE buffer washers. Bound chromatin was eluted using 250ul ChIP elution buffer (1% SDS, 100mM NaHCO3) and reverse-crosslinked. The eluates were treated with RNase A and proteinase K, and DNA was extracted via phenol-chloroform extraction and ethanol precipitation. Sequencing was carried out at Novogene.

### ChIP-seq analysis

The quality of FASTQ files (raw reads) were checked using FastQC (version. 0.11.9) and adapters were removed with fastp (*56*). Trimmed FASTQ files were aligned to the *Drosophila* genome (dm6) using Bowtie2 to generate bam files (*57*). Unmapped and low-quality reads were discarded from bam files (<=20 mapQuality) using BamTools (*58*). Duplicate reads were identified and removed from mapped reads using Picard MarkDuplicates (http://broadinstitute.github.io/picard/). Deeptools MultiBamSummary was used to determine reproducibility of ChIP-seq reads. MACS2 was used to call peaks against pooled Input/control using default settings except narrowPeaks were called for PARP-1 and broadPeaks were called for H3K4me1 (gapped), H3K9me1, H4K20me1, H3K36me1, H3K9me2, H3K9me3. Peaks were annotated to genomic features with ChIPseeker (*59*). Pairwise correlation of peaks was determined using Intervene (*60*). MACS2 bedGraph pileups were used to generate normalized coverage of ChIP-seq signals using Deeptools bigWigCompare by computing the ratio of the signals (IP vs Control/Input) using a 50bp bin size. Deeptools multiBigwigSummary was used to determine genome-wide signal correlation using a 10kb bin size. Deeptools plotHeatmap was used to create gene-centric enrichment profiles using scaled region mode. Gene clusters were determined via K means function (n=3) using Deeptools suite (*61*).

To compare H4K20me1 and PARP-1 occupancy in Human K562 cells (Supplemental Fig.S7), CrossMap was used to convert H4K20me1 bigwig from Hg19 to Hg38 to match PARP-1 bigwig. Deeptools were used to generate and plot enrichment of PARP-1 and H4K20me1 at PARP-1 gene clusters.

### Mononucleosome chromatin immunoprecipitation

We collected 60 WT wandering *Drosophila* third-instar larvae and rinsed them in 1X PBS twice. For heat shock experiments, larvae were heat-shocked in a 1.5ml DNA LoBind tube for 15mins and 30mins in a water bath at 37°C. Larvae were then homogenized using a pellet pestle homogenizer in 500ul ice-cold buffer A1 (60 mM KCl, 15 mM NaCl, 4 mM MgCl2, 15 mM 4-(2-Hydroxyethyl)piperazine-1-ethanesulfonic acid (HEPES) (pH 7.6), 0.5% Triton X-100, 0.5 mM DTT, 1×EDTA (Ethylene Diamine Tetraacetic Acid)-free protease inhibitor cocktail (Roche 04693132001). The larvae were then crosslinked by adding formaldehyde to a 1.8% concentration and incubating for 5mins at RT on a rotator. Crosslinking was stopped by adding glycine (final concentration was 0.125 M) at room temperature for 5 min on a rotator. The mixture was centrifuged at 2000g for 5 min at 4°C, and the supernatant was discarded. The pellet was washed as follows: once with 500μl A1 buffer, and once with 500μl A2 buffer (140 mM NaCl, 15 mM HEPES (pH 7.6), 1 mM EDTA, 0.5 mM ethylene glycol tetraacetic acid (EGTA), 1% Triton X-100, 0.1% sodium deoxycholate, 0.5 mM DTT, 1×EDTA-free protease inhibitor cocktail (Roche 04693132001)). For each wash, the tube was shaken for 1 min and centrifuged as before, and the supernatant was discarded. The pellet was resuspended in 500μl A2 buffer + 0.1% sodium dodecyl sulfate (SDS) and incubated on a rotating wheel at 4°C for 10 min. Then the mixture was centrifuged at 16 000g for 5 min at 4°C and the supernatant was discarded. The pellet was washed with 500μl MNase digestion buffer (10 mM Tris–HCl (pH 7.5), 15 mM NaCl, 60 mM KCl, 1 mM CaCl2, 0.15 mM spermine, 0.5 mM spermidine, 1×EDTA-free protease inhibitor cocktail) and centrifuged at 16 000g for 10 min at 4°C. The nuclei were resuspended in 500μl MNase digestion buffer and 400 U MNase (Worthington, LS004797) was added and incubated at 37°C for 20 min. The MNase digestion was terminated on ice by adding EDTA to a final concentration of 10 mM and kept on ice for 10 min. The mixture was centrifuged at 16 000g for 10 min at 4°C. The supernatant was discarded. The pellet was resuspended in 500μl A3 buffer (140 mM NaCl, 15 mM HEPES (pH 7.6), 1 mM EDTA, 0.5 mM EGTA, 1% Triton X-100, 0.1% sodium deoxycholate, 0.1% SDS, 1×EDTA-free protease inhibitor cocktail) and incubated for 30 min on a rotator at 4°C. The mixture was then centrifuged at 16 000g for 15 min at 4°C. The supernatant containing mononucleosomal fragments was transferred to a new DNA LoBind tube, then 20ul of mixture was used to check fragment size prior to immunoprecipitation. The mixture was then made up to 2ml with IP buffer (0.5% SDS, 20mM Tris, pH 8.0, 2mM EDTA, 0.5mM EGTA, 0.5mM PMSF, protease inhibitor cocktail). The mononucleosomal chromatin was pre-cleared and 800ul was incubated with 5ul anti-H4K20me1 antibody (abcam, 9051) or 5ul Rabbit IgG control at 4°C overnight and 80ul was kept as input. The immunoprecipitated chromatin was then collected with pre-washed Protein A agarose beads for 2 hours. The beads were sequentially washed with the following buffers: 1 low salt buffer wash (0.1% SDS, 1% Triton X-100, 2mM EDTA, 20mM Tris-HCL pH 8.0, 150mM NaCl), 3 high salt buffer washes (0.1% SDS, 1% Triton X-100, 2mM EDTA, 20mM Tris-HCL pH 8.0, 500mM NaCl), 1 LiCL wash (2mM EDTA, 20mM Tris-HCl pH 8.0, 0.25M LiCl, 1% NP-40) and 2 TE buffer washers before elution. Bound chromatin on beads was eluted twice at room temperature using 250ul of freshly prepared ChIP elution buffer (1% SDS, 100mM NaHCO3) for 15 minutes and reverse-crosslinked overnight. The eluates were then treated with RNase A and proteinase K prior to DNA extraction via phenol-chloroform extraction and ethanol precipitation.

### RNA sequencing

RNA was isolated from 10 wandering third-instar larvae (3 biological replicates per genotype; WT, *pr-set7^20^* and *parp-1^C03256^*) using RNeasy lipid tissue mini kit (Qiagen). RNA samples were flash-frozen in liquid nitrogen and sent to Novogene for library preparation and sequencing. mRNA was purified from total RNA via poly-T oligo beads. Libraries were prepared using Ultra II RNA library kit (NEB) and samples were sequenced on the NovaSeq 6000 platform (Illumina) at Novogene.

### RNA-seq analysis

Paired-end reads were quality-checked using FastQC and reads were mapped to the *Drosophila* genome (dm6) using RNA STAR (*62*). Reads per annotated gene was counted using featureCounts (*63*). Differential expression analysis was performed with DESeq2 (*64*), with Log2 fold change of at least 1 (absolute) considered significant (FDR < 0.05).

### Quantitative RT-PCR

For heat shock experiments, 10 third-instar larvae were heat-shocked in 1.5ml tube in a water bath at 37°C for 30 min. Total RNA was then isolated for three biological replicate using an RNeasy Lipid Tissue Mini kit (Qiagen). DNA contamination was removed using TURBO DNA-free (Ambion). RNA concentration was measured using an Agilent 2100 BioAnalyzer (Agilent Technologies) in combination with an RNA 6000 Nano LabChip (Agilent Technologies). Real-time PCR assays were run using StepOnePlus Real-Time PCR System (Applied Biosystems).

Primers used for *hsp70* ChIP-qPCR were gotten from (*19, 65*). Primers used for *hsp22*, *hsp23*, *hsp68* and *hsp83* qPCR were gotten from (*64*). Primer sequences are listed in **data file S2**.

### *pr-set7^20^* mutants validation using qRT-PCR

Total RNA was isolated from 10 third-instar larvae for three biological replicates using an RNeasy Lipid Tissue Mini kit (Qiagen). DNA contamination was removed using TURBO DNA-free (Ambion). RNA concentration was measured using an Agilent 2100 BioAnalyzer (Agilent Technologies) in combination with an RNA 6000 Nano LabChip (Agilent Technologies). Real-time PCR assays were run using StepOnePlus Real-Time PCR System (Applied Biosystems). The following primers were used: *pr-set7* (Forward) CGCACAATAGGAGTTCCC (Reverse) CCTCATCGTCCAGTTTCAG and *rp49* (Forward) GCTAAGCTGTCGCACAAAT (Reverse) GAACTTCTTGAATCCGGTGGG.

### PARP-1-EYFP re-localization in chromatin during heat shock

Larvae were grown at 20°C. Heat shock was performed by placing third instar wandering larvae individually in a 1.5ml Eppendorf tube with a small hole in the cap to let the larvae breath. The tube was placed in a Marry bath at 37°C for 30 minutes. After heat shock, the larvae were briefly washed in 1X PBS then immediately dissected in 1X supplemented Grace’s insect media (Gibco). Salivary glands were mounted in 1X supplemented Grace’s insect media (Gibco) containing Draq-5 DNA dye (Life technologies) at a 1/5000 dilution.

### Western Blot Analysis

The following antibodies were used for immunoblotting assays: anti-PR-SET7 (Rabbit, 1:1000, Novus Biologicals, 44710002), anti-H4K20me1 (Rabbit, 1:1000, Abcam, ab9051), anti-H4 (Mouse, 1:1000, Abcam, ab31830), anti-pADPr (Mouse monoclonal, 1:500, 10H - sc-56198, Santa Cruz), anti-H3 (Rabbit polyclonal, 1/1000, FL-136 sc-10809 Santa Cruz).

### Gene ontology (GO)

Gene ontology terms were determined using g:profiler (FDR<0.05) (*66*).

### Statistical analysis

Results were analyzed using the indicated statistical test in GraphPad Prism (9.4.0). Statistical significance of ChIP-seq peaks were determined using MACS2. Q-value cutoffs for RNA-seq and GO analysis were determined with DESeq2 and G:profiler respectively.

## ACKNOWLEDGEMENT

We thank Dr. Ruth Steward for sending *pr-set7^20^* null mutants. We thank Yaroslava Karpova who made the schemes of Fig.1A and 5C.

## Author contributions

G.B, G.B and N.L performed the experiments. G.B and G.B analyzed the data. G.B and A.T wrote the manuscript. G.B, G.B and A.T reviewed and edited the manuscript. G.B and A.T supervised the project.

## Data and materials availability

All data generated or analyzed during this study are included in this article. No custom code was generated in this study. ChIP-seq and RNA-seq data generated in this study are available on the GEO database (Accession no: GSE217730 and GSE222877). The following public *Drosophila* third-instar larvae ChIP-seq (ENCODE) were used in this study: GSE47282 (H3K4me1), GSE47254 (H4K20me1), GSE47249 (H3K36me1), GSE47289 (H3K9me1), GSE47260 (H3K9me2) and GSE47258 (H3K9me3), GSE15292 (RNA-Polymerase II). In addition, we used the Developmental time-course RNA-seq dataset (*67*), SRP001065. Finaly, the following public Human K562 cells ChIP-seq (ENCODE) were used in this study: GSE206022 (PARP-1 CUT&Tag), GSE29611 (H4K20me1).

## REFERENCES

1. Kouzarides T. Chromatin modifications and their function. Cell. 2007;128(4):693–705.

2. Peterson CL, Laniel MA. Histones and histone modifications. Curr Biol. 2004;14(14):R546–51.

3. Kraus WL, Hottiger MO. PARP-1 and gene regulation: progress and puzzles. Mol Aspects Med. 2013;34(6):1109–23.

4. Posavec Marjanovic M, Crawford K, Ahel I. PARP, transcription and chromatin modeling. Semin Cell Dev Biol. 2017;63:102–13.

5. Thomas C, Tulin AV. Poly-ADP-ribose polymerase: machinery for nuclear processes. Mol Aspects Med. 2013;34(6):1124–37.

6. D’Amours D, Desnoyers S, D’Silva I, Poirier GG. Poly(ADP-ribosyl)ation reactions in the regulation of nuclear functions. Biochem J. 1999;342 ( Pt 2)(Pt 2):249–68.

7. Tulin A, Spradling A. Chromatin loosening by poly(ADP)-ribose polymerase (PARP) at Drosophila puff loci. Science. 2003;299(5606):560-2.

8. Berger SL. Histone modifications in transcriptional regulation. Curr Opin Genet Dev. 2002;12(2):142–8.

9. Vaquero A, Loyola A, Reinberg D. The constantly changing face of chromatin. Sci Aging Knowledge Environ. 2003;2003(14):RE4.

10. Bannister AJ, Kouzarides T. Regulation of chromatin by histone modifications. Cell Res. 2011;21(3):381–95.

11. Kotova E, Lodhi N, Jarnik M, Pinnola AD, Ji Y, Tulin AV. Drosophila histone H2A variant (H2Av) controls poly(ADP-ribose) polymerase 1 (PARP1) activation in chromatin. Proceedings of the National Academy of Sciences. 2011;108(15):6205–10.

12. Pinnola A, Naumova N, Shah M, Tulin AV. Nucleosomal core histones mediate dynamic regulation of poly(ADP-ribose) polymerase 1 protein binding to chromatin and induction of its enzymatic activity. J Biol Chem. 2007;282(44):32511–9.

13. Barski A, Cuddapah S, Cui K, Roh TY, Schones DE, Wang Z, et al. High-resolution profiling of histone methylations in the human genome. Cell. 2007;129(4):823–37.

14. Karachentsev D, Sarma K, Reinberg D, Steward R. PR-Set7-dependent methylation of histone H4 Lys 20 functions in repression of gene expression and is essential for mitosis. Genes Dev. 2005;19(4):431–5.

15. Kotova E, Jarnik M, Tulin AV. Uncoupling of the transactivation and transrepression functions of PARP1 protein. Proc Natl Acad Sci U S A. 2010;107(14):6406–11.

16. Esteve PO, Sen S, Vishnu US, Ruse C, Chin HG, Pradhan S. Poly ADP-ribosylation of SET8 leads to aberrant H4K20 methylation in mammalian nuclear genome. Commun Biol. 2022;5(1):1292.

17. Petesch SJ, Lis JT. Activator-induced spread of poly(ADP-ribose) polymerase promotes nucleosome loss at Hsp70. Mol Cell. 2012;45(1):64–74.

18. Thomas C, Ji Y, Wu C, Datz H, Boyle C, MacLeod B, et al. Hit and run versus long-term activation of PARP-1 by its different domains fine-tunes nuclear processes. Proc Natl Acad Sci U S A. 2019;116(20):9941–6.

19. Petesch SJ, Lis JT. Rapid, transcription-independent loss of nucleosomes over a large chromatin domain at Hsp70 loci. Cell. 2008;134(1):74–84.

20. Oda H, Hubner MR, Beck DB, Vermeulen M, Hurwitz J, Spector DL, et al. Regulation of the histone H4 monomethylase PR-Set7 by CRL4(Cdt2)-mediated PCNA-dependent degradation during DNA damage. Mol Cell. 2010;40(3):364–76.

21. Wang Y, Jia S. Degrees make all the difference: the multifunctionality of histone H4 lysine 20 methylation. Epigenetics. 2009;4(5):273–6.

22. De Angelis Campos AC, Rodrigues MA, de Andrade C, de Goes AM, Nathanson MH, Gomes DA. Epidermal growth factor receptors destined for the nucleus are internalized via a clathrin-dependent pathway. Biochem Biophys Res Commun. 2011;412(2):341–6.

23. Gibson BA, Zhang Y, Jiang H, Hussey KM, Shrimp JH, Lin H, et al. Chemical genetic discovery of PARP targets reveals a role for PARP-1 in transcription elongation. Science. 2016;353(6294):45–50.

24. Krishnakumar R, Kraus WL. PARP-1 regulates chromatin structure and transcription through a KDM5B-dependent pathway. Mol Cell. 2010;39(5):736–49.

25. Nalabothula N, Al-jumaily T, Eteleeb AM, Flight RM, Xiaorong S, Moseley H, et al. Genome-Wide Profiling of PARP1 Reveals an Interplay with Gene Regulatory Regions and DNA Methylation. PLoS One. 2015;10(8):e0135410.

26. Nikolaou KC, Moulos P, Harokopos V, Chalepakis G, Talianidis I. Kmt5a Controls Hepatic Metabolic Pathways by Facilitating RNA Pol II Release from Promoter-Proximal Regions. Cell Rep. 2017;20(4):909–22.

27. Talasz H, Lindner HH, Sarg B, Helliger W. Histone H4-lysine 20 monomethylation is increased in promoter and coding regions of active genes and correlates with hyperacetylation. J Biol Chem. 2005;280(46):38814–22.

28. Vakoc CR, Sachdeva MM, Wang H, Blobel GA. Profile of histone lysine methylation across transcribed mammalian chromatin. Mol Cell Biol. 2006;26(24):9185–95.

29. Beck DB, Oda H, Shen SS, Reinberg D. PR-Set7 and H4K20me1: at the crossroads of genome integrity, cell cycle, chromosome condensation, and transcription. Genes Dev. 2012;26(4):325–37.

30. Ji Y, Tulin AV. The roles of PARP1 in gene control and cell differentiation. Curr Opin Genet Dev. 2010;20(5):512–8.

31. Khoury-Haddad H, Guttmann-Raviv N, Ipenberg I, Huggins D, Jeyasekharan AD, Ayoub N. PARP1-dependent recruitment of KDM4D histone demethylase to DNA damage sites promotes double-strand break repair. Proc Natl Acad Sci U S A. 2014;111(7):E728–37.

32. Tuzon CT, Spektor T, Kong X, Congdon LM, Wu S, Schotta G, et al. Concerted activities of distinct H4K20 methyltransferases at DNA double-strand breaks regulate 53BP1 nucleation and NHEJ-directed repair. Cell Rep. 2014;8(2):430–8.

33. Yang L, Huang K, Li X, Du M, Kang X, Luo X, et al. Identification of poly(ADP-ribose) polymerase-1 as a cell cycle regulator through modulating Sp1 mediated transcription in human hepatoma cells. PLoS One. 2013;8(12):e82872.

34. Bamgbose G, Tulin A. PARP-1 is a transcriptional rheostat of metabolic and bivalent genes during development. Life Sci Alliance. 2024;7(2).

35. Bamgbose G, Johnson S, Tulin A. Cooperative targeting of PARP-1 domains to regulate metabolic and developmental genes. Front Endocrinol (Lausanne). 2023;14:1152570.

36. Arbeitman MN, Furlong EE, Imam F, Johnson E, Null BH, Baker BS, et al. Gene expression during the life cycle of Drosophila melanogaster. Science. 2002;297(5590):2270-5.

37. White KP, Rifkin SA, Hurban P, Hogness DS. Microarray analysis of Drosophila development during metamorphosis. Science. 1999;286(5447):2179-84.

38. Nishimura T. Feedforward Regulation of Glucose Metabolism by Steroid Hormones Drives a Developmental Transition in Drosophila. Curr Biol. 2020;30(18):3624–32 e5.

39. Tanaka H, Takebayashi SI, Sakamoto A, Igata T, Nakatsu Y, Saitoh N, et al. The SETD8/PR-Set7 Methyltransferase Functions as a Barrier to Prevent Senescence-Associated Metabolic Remodeling. Cell Rep. 2017;18(9):2148–61.

40. Veloso A, Kirkconnell KS, Magnuson B, Biewen B, Paulsen MT, Wilson TE, et al. Rate of elongation by RNA polymerase II is associated with specific gene features and epigenetic modifications. Genome Res. 2014;24(6):896–905.

41. Wang J, Telese F, Tan Y, Li W, Jin C, He X, et al. LSD1n is an H4K20 demethylase regulating memory formation via transcriptional elongation control. Nat Neurosci. 2015;18(9):1256–64.

42. Fuchs G, Hollander D, Voichek Y, Ast G, Oren M. Cotranscriptional histone H2B monoubiquitylation is tightly coupled with RNA polymerase II elongation rate. Genome Res. 2014;24(10):1572–83.

43. Abbas T, Shibata E, Park J, Jha S, Karnani N, Dutta A. CRL4(Cdt2) regulates cell proliferation and histone gene expression by targeting PR-Set7/Set8 for degradation. Mol Cell. 2010;40(1):9–21.

44. Kapoor-Vazirani P, Vertino PM. A dual role for the histone methyltransferase PR-SET7/SETD8 and histone H4 lysine 20 monomethylation in the local regulation of RNA polymerase II pausing. J Biol Chem. 2014;289(11):7425–37.

45. Shoaib M, Chen Q, Shi X, Nair N, Prasanna C, Yang R, et al. Histone H4 lysine 20 mono-methylation directly facilitates chromatin openness and promotes transcription of housekeeping genes. Nat Commun. 2021;12(1):4800.

46. Gungi A, Saha S, Pal M, Galande S. H4K20me1 plays a dual role in transcriptional regulation of regeneration and axis patterning in Hydra. Life Sci Alliance. 2023;6(5).

47. Shi X, Kachirskaia I, Yamaguchi H, West LE, Wen H, Wang EW, et al. Modulation of p53 function by SET8-mediated methylation at lysine 382. Mol Cell. 2007;27(4):636–46.

48. Takawa M, Cho HS, Hayami S, Toyokawa G, Kogure M, Yamane Y, et al. Histone lysine methyltransferase SETD8 promotes carcinogenesis by deregulating PCNA expression. Cancer Res. 2012;72(13):3217–27.

49. Rickels R, Herz HM, Sze CC, Cao K, Morgan MA, Collings CK, et al. Histone H3K4 monomethylation catalyzed by Trr and mammalian COMPASS-like proteins at enhancers is dispensable for development and viability. Nat Genet. 2017;49(11):1647–53.

50. Chory EJ, Calarco JP, Hathaway NA, Bell O, Neel DS, Crabtree GR. Nucleosome Turnover Regulates Histone Methylation Patterns over the Genome. Mol Cell. 2019;73(1):61–72 e3.

51. Liu Z, Kraus WL. Catalytic-Independent Functions of PARP-1 Determine Sox2 Pioneer Activity at Intractable Genomic Loci. Mol Cell. 2017;65(4):589–603 e9.

52. Tulin A, Stewart D, Spradling AC. The Drosophila heterochromatic gene encoding poly(ADP-ribose) polymerase (PARP) is required to modulate chromatin structure during development. Genes Dev. 2002;16(16):2108–19.

53. Muthurajan UM, Hepler MR, Hieb AR, Clark NJ, Kramer M, Yao T, et al. Automodification switches PARP-1 function from chromatin architectural protein to histone chaperone. Proc Natl Acad Sci U S A. 2014;111(35):12752–7.

54. Artavanis-Tsakonas S. Accessing the Exelixis collection. Nat Genet. 2004;36(3):207.

55. Johansen KM, Cai W, Deng H, Bao X, Zhang W, Girton J, et al. Polytene chromosome squash methods for studying transcription and epigenetic chromatin modification in Drosophila using antibodies. Methods. 2009;48(4):387–97.

56. Chen S, Zhou Y, Chen Y, Gu J. fastp: an ultra-fast all-in-one FASTQ preprocessor. Bioinformatics. 2018;34(17):i884–i90.

57. Langmead B, Salzberg SL. Fast gapped-read alignment with Bowtie 2. Nat Methods. 2012;9(4):357–9.

58. Barnett DW, Garrison EK, Quinlan AR, Stromberg MP, Marth GT. BamTools: a C++ API and toolkit for analyzing and managing BAM files. Bioinformatics. 2011;27(12):1691–2.

59. Yu G, Wang LG, He QY. ChIPseeker: an R/Bioconductor package for ChIP peak annotation, comparison and visualization. Bioinformatics. 2015;31(14):2382–3.

60. Khan A, Mathelier A. Intervene: a tool for intersection and visualization of multiple gene or genomic region sets. BMC Bioinformatics. 2017;18(1):287.

61. Ramirez F, Ryan DP, Gruning B, Bhardwaj V, Kilpert F, Richter AS, et al. deepTools2: a next generation web server for deep-sequencing data analysis. Nucleic Acids Res. 2016;44(W1):W160–5.

62. Dobin A, Davis CA, Schlesinger F, Drenkow J, Zaleski C, Jha S, et al. STAR: ultrafast universal RNA-seq aligner. Bioinformatics. 2013;29(1):15–21.

63. Liao Y, Smyth GK, Shi W. featureCounts: an efficient general purpose program for assigning sequence reads to genomic features. Bioinformatics. 2014;30(7):923–30.

64. Love MI, Huber W, Anders S. Moderated estimation of fold change and dispersion for RNA-seq data with DESeq2. Genome Biol. 2014;15(12):550.

65. Berson A, Sartoris A, Nativio R, Van Deerlin V, Toledo JB, Porta S, et al. TDP-43 Promotes Neurodegeneration by Impairing Chromatin Remodeling. Curr Biol. 2017;27(23):3579–90 e6.

66. Raudvere U, Kolberg L, Kuzmin I, Arak T, Adler P, Peterson H, et al. g:Profiler: a web server for functional enrichment analysis and conversions of gene lists (2019 update). Nucleic Acids Res. 2019;47(W1):W191–W8.

67. Graveley BR, Brooks AN, Carlson JW, Duff MO, Landolin JM, Yang L, et al. The developmental transcriptome of Drosophila melanogaster. Nature. 2011;471(7339):473–9.

