## Supplementary material for "Mono-methylated histones control PARP-1 in chromatin and transcription": Supplememtal data

**THIS FILE INCLUDES:**

**Supplemental Figure S1.** PARP-1 binding is inhibited by H3K9me2/3 peptides or by phosphorylation of adjacent residues.

**Supplemental Figure S2.** Inhibition of PARP-1 binding by H3K9me2/3 and phosphorylation of adjacent residues.

**Supplemental Figure S3.** Validation of *pr-set7*$^{20}$ mutant.

**Supplemental Fig.S4.** PR-SET7 RNA level is not affected in *parp-1*$^{C03256}$ mutant.

**Supplemental Fig.S5.** PR-SET7 protein level is not affected in *parp-1*$^{C03256}$ mutant.

**Supplemental Fig.S6.** Genes coenriched in H4K20me1 and PARP-1 that are upregulated in both *parp-1* and *pr-set7* mutants are highly expressed genes.

**Supplemental Fig.S7.** PARP-1 binding correlates with H4K20me1 enrichment in Human K562 cells.

**Data file S1 (Separate file).** PARP-1_histone_peptide_array_results.

**Data file S2 (Separate file).** Primers used in this study.

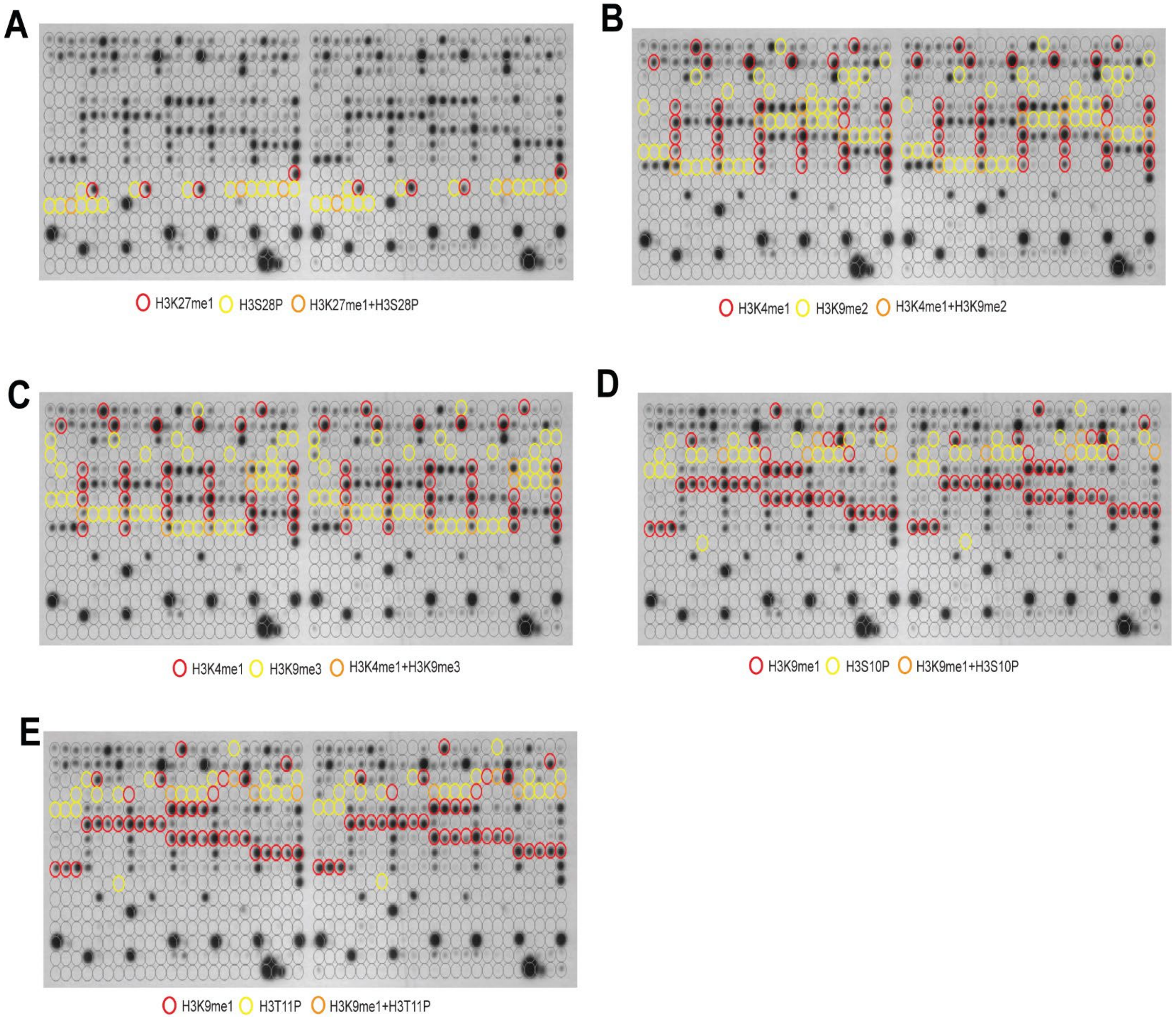


**Supplemental Fig.S1. PARP-1 binding is inhibited by H3K9me2/3 peptides or by phosphorylation of adjacent residues.** Signal intensity on modified histone peptide array showing inhibition of PARP-1 binding to spots containing H3K27me1 by the presence of H3S28P peptides (**A**), inhibition of PARP-1 binding to spots containing H3K4me1 peptides by the presence of H3K9me2 (**B**) or H3K9me3 (**C**) peptides, inhibition of PARP-1 binding to spots containing H3K9me1 peptides by the presence of H3S10P (**D**) or H3T11P (**E**) peptides.

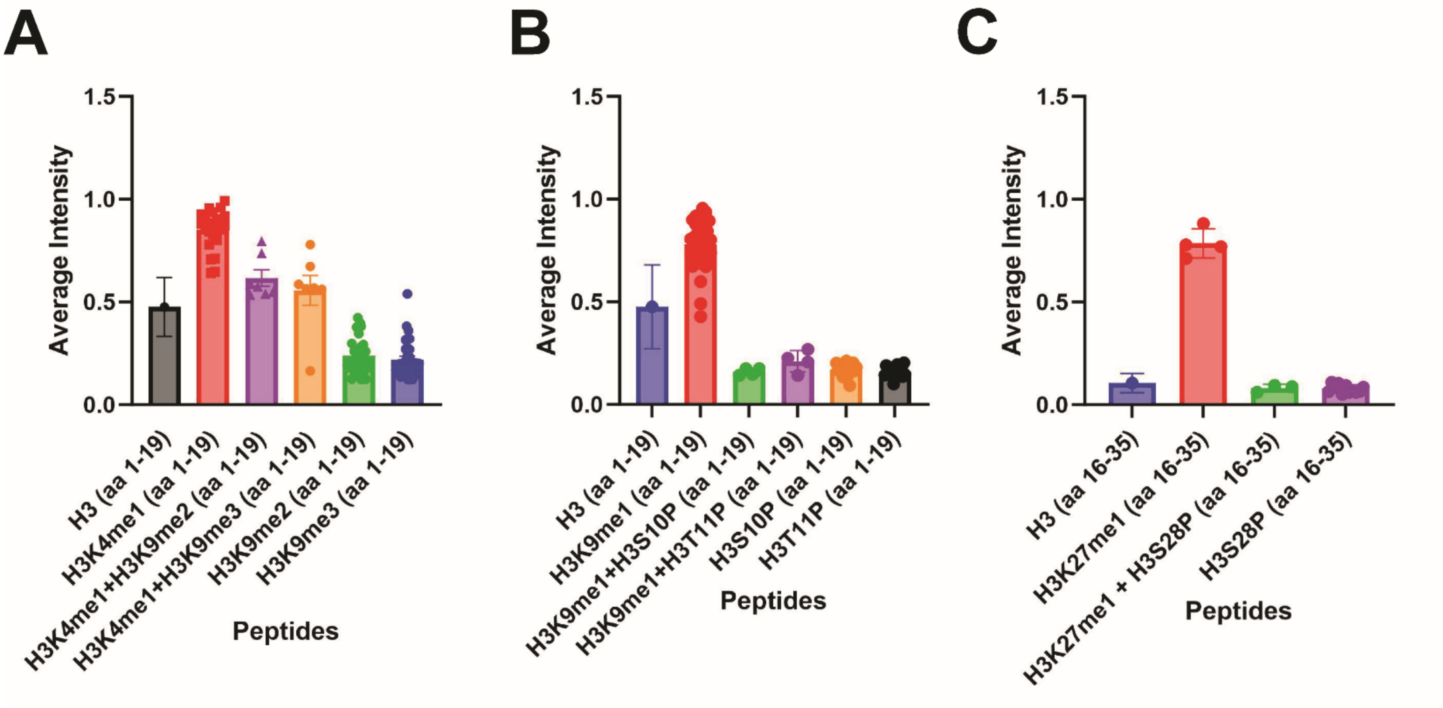


**Supplemental Fig.S2. Inhibition of PARP-1 binding by H3K9me2/3 and phosphorylation of adjacent residues.** Average intensities from histone peptide array showing the inhibition of PARP-1 binding to spots containing H3K4me1 peptides by the presence of H3K9me2/3 peptide (**A**), inhibition of PARP-1 binding to spots containing H3K9me1 peptides by the presence of H3S10P or H3T11P peptides (**B**), inhibition of PARP-1 binding to spots containing H3K27me1 peptides by the presence of H3S28P peptides (**C**).

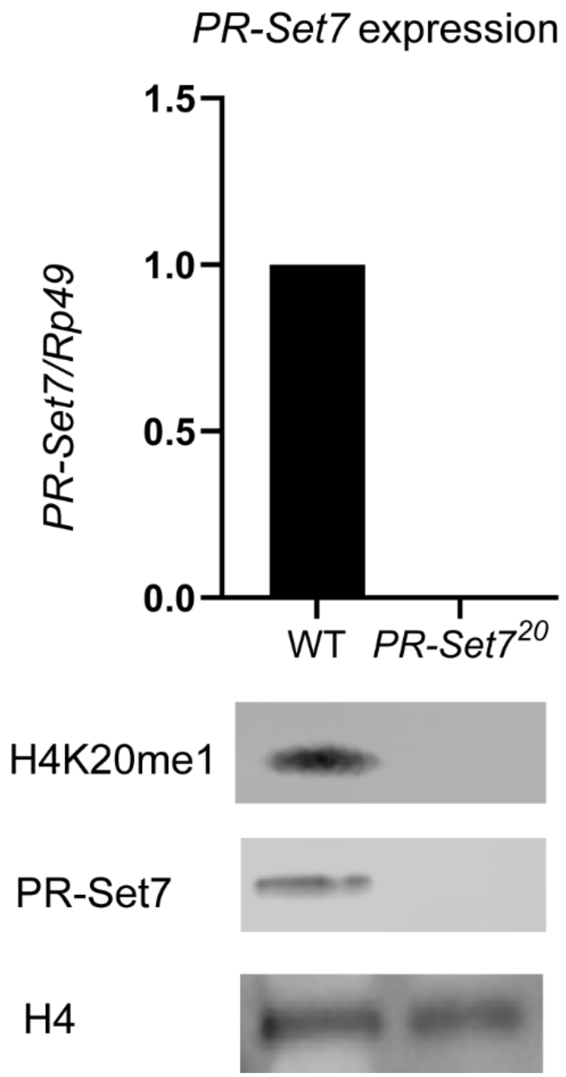


**Supplemental Fig.S3. Validation of *pr-set7*$^{20}$ mutant. (Top)** Expression levels of PR-SET7 in WT or *pr-set7*$^{20}$ mutant during third-instar larval stage (3 biological replicates). (**Bottom**) Western blot of H4K20me1, PR-SET7 and H4 loading control in WT or *pr-set7*$^{20}$ mutant during third-instar larval stage (2 biological replicates).

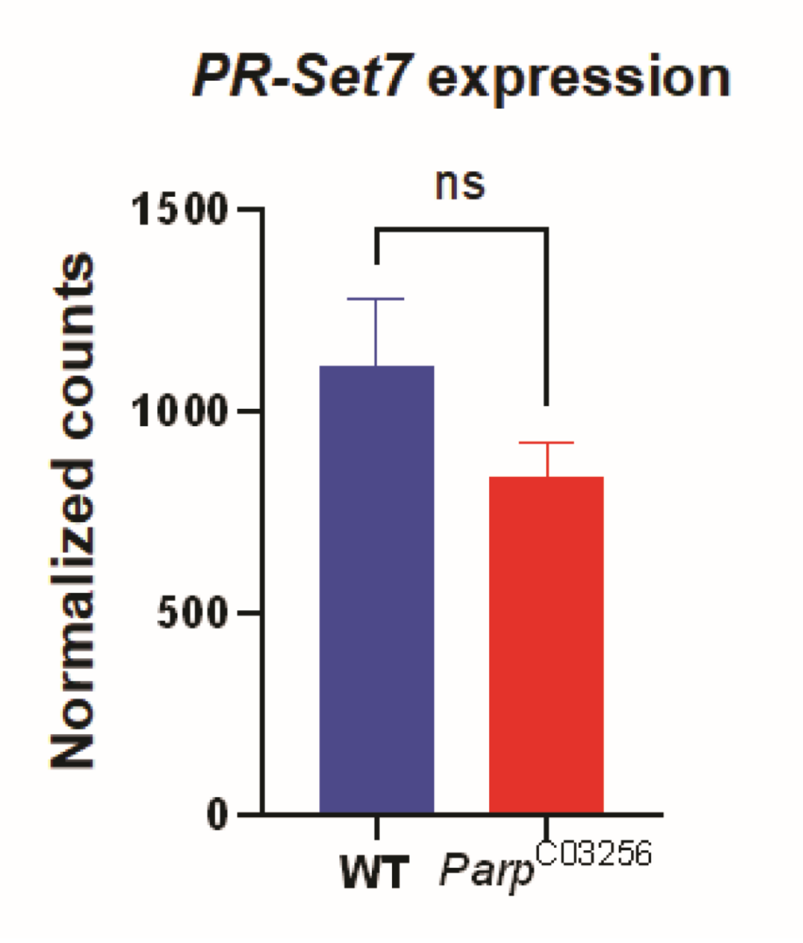


**Supplemental Fig.S4. PR-SET7 expression level is not affected in *parp-1*$^{C03256}$ mutant.** Normalized counts of Pr-SET7 RNA in a Wildtype (blue) or in a *parp-1*$^{C03256}$ (red) background based on RNA-seq data (GSE222877). The experiment was performed in triplicates. The statistical test is an unpaired two-tailed *t*-test. **ns**: non-significant.

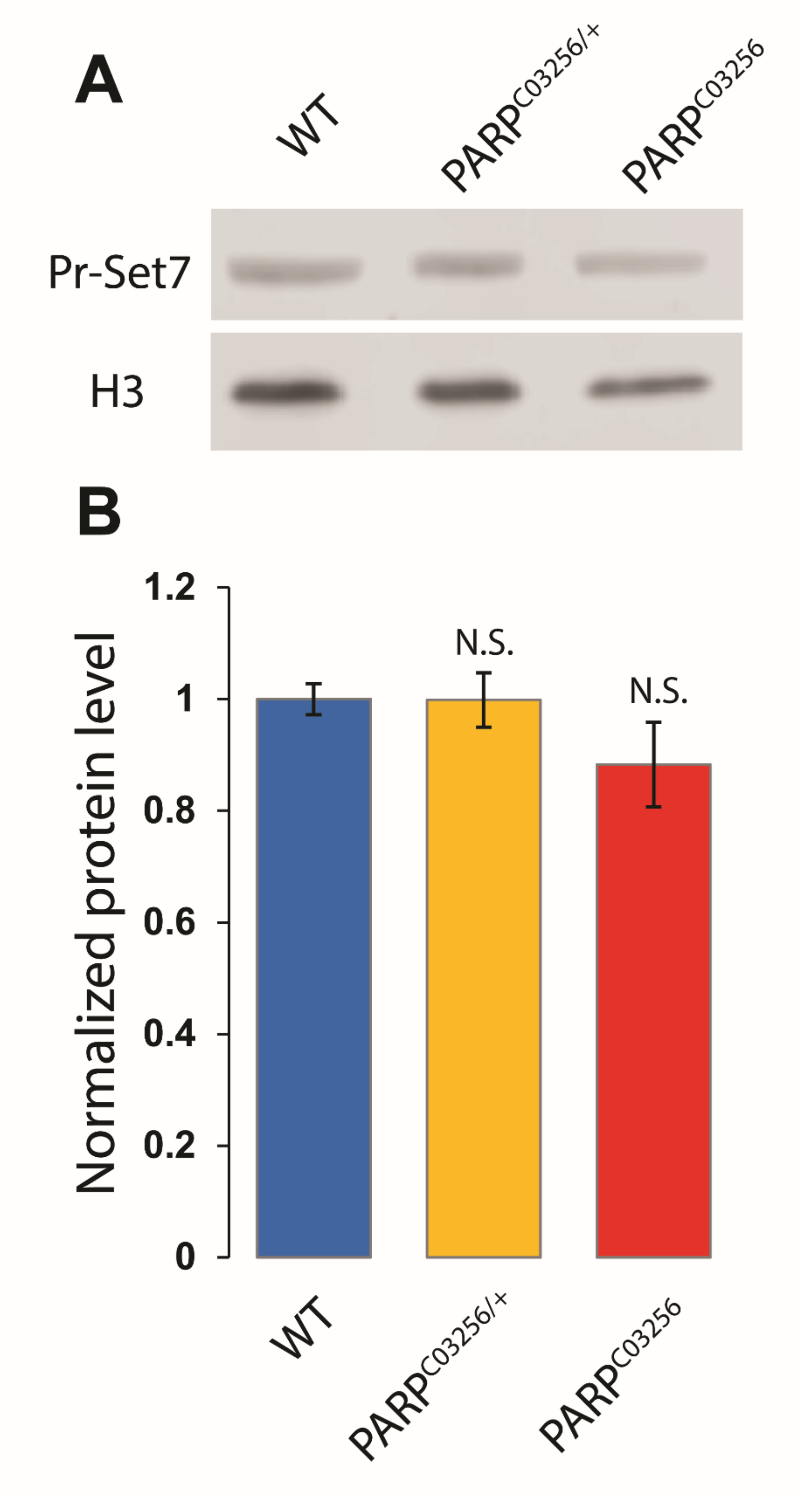


**Supplemental Fig.S5. PR-SET7 protein level is not affected in *parp-1*$^{C03256}$ mutant. A.** Western blot showing Pr-SET7 protein level in a wild type (WT), *parp-1* heterozygote (*Parp*$^{C03256/+}$), or homozygote mutant (*Parp*$^{C03256}$) background. H3 is used as a loading control. **B.** Quantification of PR-SET7 protein level based on three independent biological replicates and normalized to H3 level. The statistical test is an unpaired two-tailed *t*-test. N.S: non-significant.

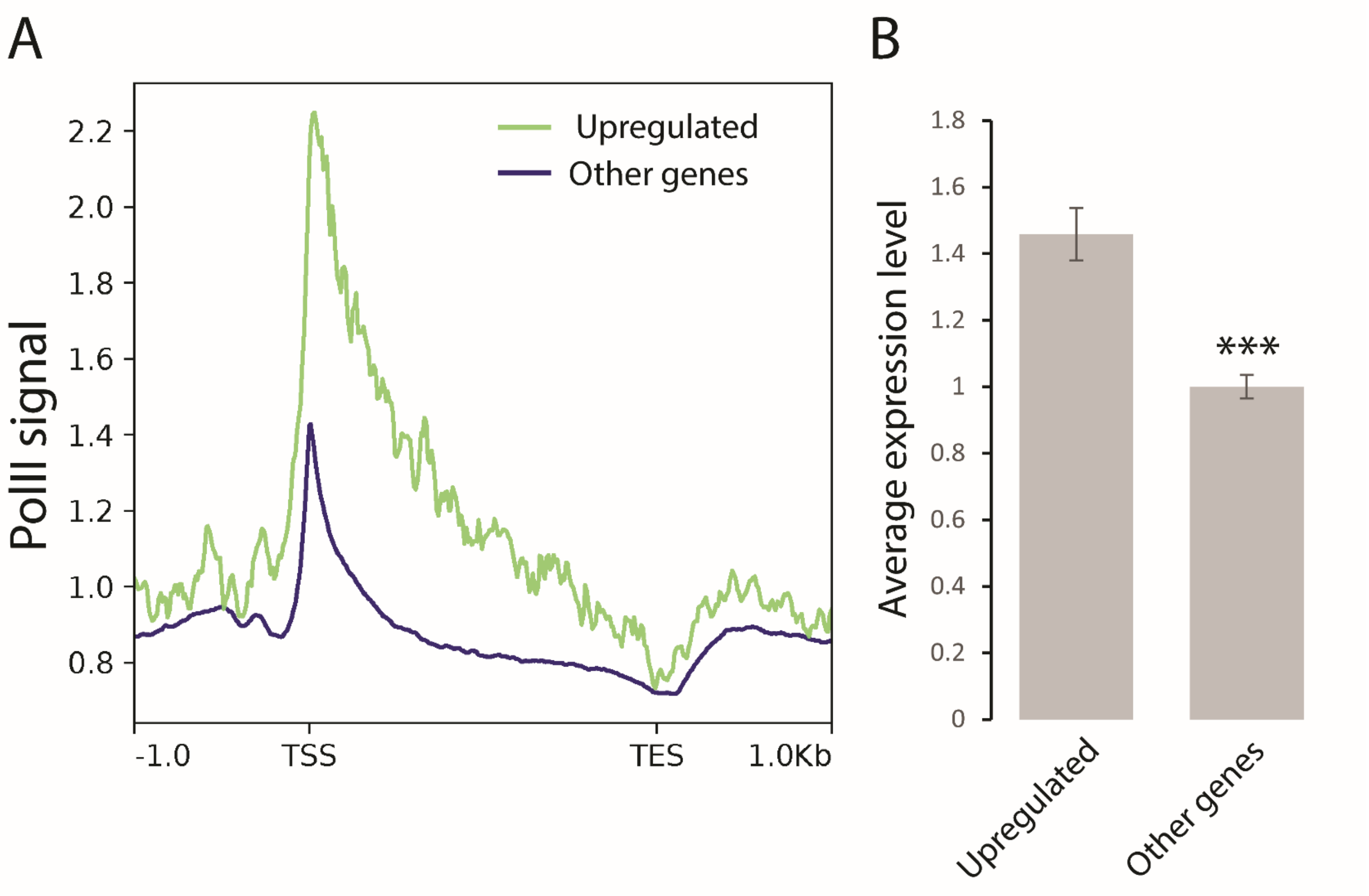


**Supplemental Fig.S6. Genes coenriched in H4K20me1 and PARP-1 that are upregulated in both *parp-1* and *pr-set7* mutants are highly expressed genes. A.** Distribution of RNA-Polymerase II (PolII) protein along genes that coenriched by H4K20me1 and PARP-1 and upregulated both *parp-1* and *pr-set7* mutants (green) and along all other *Drosophila* genes (blue). The region of all target genes is scaled to a 2kb region (from TSS to TES) and includes the 1kb region upstream from TSS and downstream from TES. The analysis was performed on the whole organism on *Drosophila* third instar larval puffstage 7-9. **B.** Average expression level of genes coenriched by H4K20me1 and PARP-1 and upregulated both *parp-1* and *pr-set7* mutants (upregulated) and average expression of all other genes. The expression was measured on the whole organism on *Drosophila* third instar larval puffstage 7-9. Statistical test is an unpaired two-tailed *t*-test. ***: $p$-value < 0.01.

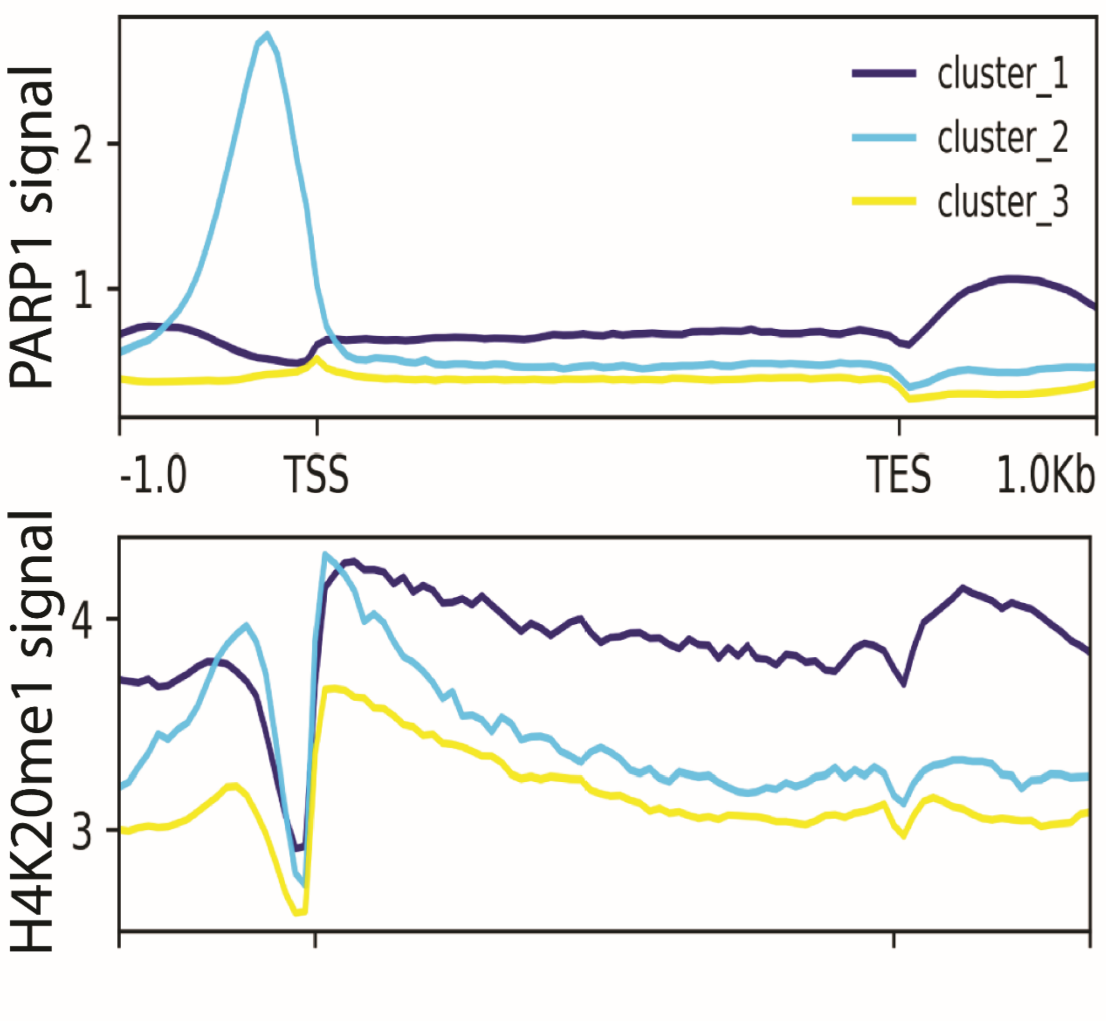


**Supplemental Fig.S7. PARP-1 binding correlates with H4K20me1 enrichment in Human K562 cells.** Metagene plots showing enrichment of PARP-1 (CUT&Tag) and H4K20me1 (ChIP-Seq) at PARP-1 clusters (cluster 1 = 5385, cluster 2 = 2637, cluster 3 = 11, 920 regions) in human protein-coding genes. Signals extend from −1 kb of the TSS to +1 kb of the TES.
